# How do the molecular determinants of antiviral resistance shape the dynamic of phytoplankton-virus interaction?

**DOI:** 10.64898/2026.09.29.755341

**Authors:** Raphaël Rousseau, Carlos Cáceres, Gwenaël Piganeau, Sébastien Gourbière

## Abstract

Phytoplankton sustain all marine ecosystems and are key components of the carbon cycle being responsible for ∼50% of the global primary production. A major hypothesis to explain their coexistence with highly efficient lytic viruses is antiviral resistance. Although intracellular and extracellular mechanisms have been shown to underly such resistance and while its determinism is commonly associated to a genomic polymorphism or a (random or virus induced) phenotypic plasticity, comparisons of the impact of those molecular determinants on phytoplankton-virus population dynamic are genuinely lacking. We combined 12 epidemiological models with an unprecedented set of parameter estimates derived from experimental studies of the ubiquitous species *Ostreococcus tauri* and its prasinoviruses to provide such comprehensive predictions. We show that ‘bet-hedging’ strategies corresponding to random phenotypic switches between susceptible and resistant cells provide the strongest barrier to virus emergence. When such resistance is associated to intracellular mechanisms the virus’ R0 is at least halved because of the strong ‘dilution effect’ produced by the ‘dead-end’ hosts. In addition, once the virus has spread, both random and virus-induced phenotypic plasticity strongly stabilize the otherwise ‘boom and bust’ phytoplankton-virus dynamics, which is highly consistent with the experimental dynamics observed for such interactions.

## Introduction

Phytoplankton are essential components at the base of the food chain that sustain marine and freshwater ecosystems (Naselli-Flores and Padisák 2023). While these autotrophic organisms account for less than 1% of the global biomass (Bar-On et al. 2018), they are key components of the major biogeochemical cycles, and especially of the carbon cycle, as they provide about half of the world primary production and atmospheric oxygen through their photosynthetic activity (Litchman et al. 2015).

The abundance and diversity of marine phytoplankton species are under the ‘top-down’ control of viruses (Brussaard 2004) that stand as the most abundant entities in oceanic waters (Fuhrman 1999; Suttle 2007). Lytic infections indeed constitute a predominant factor of phytoplankton bloom’s regulation (Brussaard 2003; Diaz et al. 2023), thereby contributing to the remobilisation of the corresponding organic matter with significant effects on the carbon cycle through the so-called ‘viral shunt’ (Wilhelm and Suttle 1999) and ‘viral shuttle’ (Weinbauer 2004; Sullivan et al. 2017). Intriguingly, while most phytoplankton populations undergo substantial lytic events (Suttle 2007), they broadly coexist with their viruses in marine environments (Baran et al. 2017; Tomaru et al. 2018), while those provide no obvious spatial or temporal refuges. A major hypothesis to resolve such an apparent paradox is the existence of antiviral resistance, which has been documented for numerous phytoplankton-virus pairs both in laboratory cultures (Thyrhaug et al. 2003; Tomaru et al. 2009; Kimura and Tomaru 2014; Zborowsky and Lindell 2019; Yau et al. 2018, 2020; Esmael et al. 2023; Bedi De Silva et al. 2024; Shaler et al. 2025) and in marine environments (Schleyer et al. 2023). While different mechanisms have been shown to underly such antiviral resistance and while they can be associated to genomic mutations or to a phenotypic plasticity (see below), there is no comprehensive understanding of the quantitative implications of these alternative mechanisms and determinisms on the dynamics of phytoplankton-virus interactions.

The mechanisms of phytoplankton resistance to their viruses have long been classified according to the stage of the infection at which they occur. First, extracellular mechanisms can block the adsorption of viral particles or their entry into unicellular hosts. This can be mediated by surface molecules that prevent the recognition of the cells, or by the production of extracellular polymeric substances (EPS) limiting contacts with the hosts. In the former case, host cells become ‘invisible’ to the virus by modification or suppression of their receptors (Avrani et al. 2011; Yau et al. 2018; Schleyer et al. 2023; Bedi De Silva et al. 2024). In the latter, the EPS-mediated resistance involves the production of exopolysaccharides providing a mucus that act as a physical barrier or a viral inhibitor (Brussaard et al. 2007; Vincent et al. 2021). Second, intracellular mechanisms can disrupt viral infection by weakening or blocking the virus replication, or through the control of its release into the marine environment (Tomaru et al. 2009; Thomas et al. 2011; Zborowsky and Lindell 2019; Esmael et al. 2023). While all the mechanisms described above primarily benefit the resistant individuals, programmed cell death (PCD) provides an alternative collective defence process to various unicellular organisms (Bidle and Falkowski 2004; Bidle et al. 2007; Bidle 2015). Even though the cells playing such an ‘altruistic suicide’ strategy do not actually resist the viral interaction, PCD can reduce the amount of environmental viral particles to which the rest of the host population is exposed. Although the actual impact of virally induced PCD on phytoplankton populations remains discussed (Bidle and Vardi 2011; Bidle 2015; Vincent et al. 2021), it is likely to act as a mechanism complementing the above alternative forms of individual resistance.

Omics analyses allowing to unravel the genomic and molecular determinisms of phytoplankton individual resistance and their variability are still emerging (Yau et al. 2020; Shaler et al. 2025; Schleyer et al. 2023; Zborowsky et al. 2025; James et al. 2026). A typical determinism underlying resistance corresponds to genomic mutations (GM) (Sasaki 2002), either punctual (point mutations) or chromosomic (structural variations), and the antiviral resistance of different phytoplankton species has indeed been associated to major structural rearrangements of chromosomes or genomic islands through genomic studies (Avrani et al. 2011; Bedi De Silva et al. 2024). Meanwhile, transcriptomic, epi-genomic and/or metabolomic analyses have provided further evidence that the regulation of genomic regions associated to phytoplankton resistance to their viruses can be the determinants of a phenotypic plasticity (PP) allowing individuals to switch between susceptible and resistant states. However, the proximate causes for such a PP, that can potentially occur randomly as a result of a ‘bet-hedging’ strategy (Yau et al. 2020) or through a more deterministic induction by the virus, remain difficult to disentangle (Thyrhaug et al. 2003; Brussaard et al. 2007; Yau et al. 2018, 2020; Shaler et al. 2025).

The experimental assessments and comparisons of the population scale impacts of the above mechanisms and determinisms of phytoplankton individual antiviral resistance represent a time-consuming and resource-intensive challenge, since it remains difficult to monitor long term host virus interactions while maintaining axenic culture (but see (Lee et al. 2021; Schiano Di Visconte et al. 2022)). This obviously impedes our understanding of their consequences on the ecological and evolutionary dynamics of phytoplankton-virus interactions in natural marine environments. Epidemiological models have proven to be essential tools to predict how host and pathogen individual life-history traits influence the dynamics of their interactions (Keeling and Rohani 2011). Although such models have been developed to assess the impact of lytic viruses on the persistence and abundance of phytoplankton (see (Middleton et al. 2017; Listmann et al. 2026) for reviews and (Fuhrman et al. 2011; Béchette et al. 2013; Edwards and Steward 2018; Demory et al. 2021; Zhang et al. 2024)), only a few of those account for antiviral resistance (Yau et al. 2020; Frémont et al. 2026). Accordingly, there is currently no comprehensive modelling to provide systematic predictions about the quantitative implications of the above alternative mechanisms and determinisms of antiviral resistance on the dynamics of unicellular phytoplankton-viruses interactions.

In this contribution, we intend to develop such a theoretical framework to allow scaling-up from the description of antiviral resistance at the individual level to the dynamics of lytic virus transmission through a phytoplankton population. We aim at modelling any pair of known mechanism and determinism of such resistance that are described above and to make comparative predictions about their potential effects on the corresponding host-virus ecological dynamics. More specifically, we intend to answer the three following questions. First, how could (the individual cost of) antiviral resistance affects the growth of a phytoplankton population in the absence of virus? Second, how does resistance reduce the rate of viral spread into such a virus-free environment? Third, once the virus has spread into a phytoplankton population, how does resistance influence the stability of their coexistence? To answer these questions, we analysed the comprehensive set of phytoplankton-virus epidemiological models that we propose, which allowed providing both general analytical results and more specific predictions. The latter were obtained by tailoring our modelling to *Ostreococcus tauri*, a key species of the *Mamiellophyceae* that corresponds to the most abundant class of green algae in coastal waters (Tragin and Vaulot 2018). These ubiquitous species have been repeatedly shown to establish antiviral resistance to limit the impact of their lytic prasinoviruses (Yau et al. 2018; Thomas et al. 2011; Thomy et al. 2026; Yau et al. 2016).

## Methods

### **The** core SIV model of phytoplankton-virus interactions

The core of our modelling is made of a ‘SIV’ representation of phytoplankton-virus interactions allowing to predict the temporal dynamics of the density of susceptible (S) and infected (I) phytoplankton cells, together with the number of virions in the environment (V) (Fig. 1a). The pool of susceptible host cells grows according to the difference between their rates of asexual reproduction (*b*) and intrinsic mortality (*d*) and is downregulated by intraspecific competition that typically limits the rate of cell division. The overall impact of such a competition was considered to rise linearly with the size of the phytoplankton population (N = S + I) and according to the strength of the interaction between individuals (*c*). Susceptible cells become infected upon contacts with virions at a rate depending on their individual encounter rate (*β*) and the densities S and V. The subsequent within-host dynamic of lytic viruses ends with the breaking down of the membrane of the infected cell at a ‘lysis’ rate (*α*) and the release of a number of virions in the environment (*b_v_*) commonly called the ‘burst-size’. Finally, the free viral particles tend to become ineffective because of deleterious environmental conditions, which was accounted for through a fixed decay rate (*d_v_*). These descriptions of the basic processes underlying the interaction dynamics between unicellular phytoplankton and their lytic viruses led to the following SIV model;

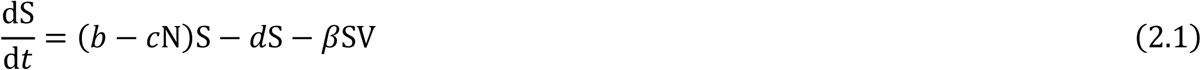

**Figure 1.**
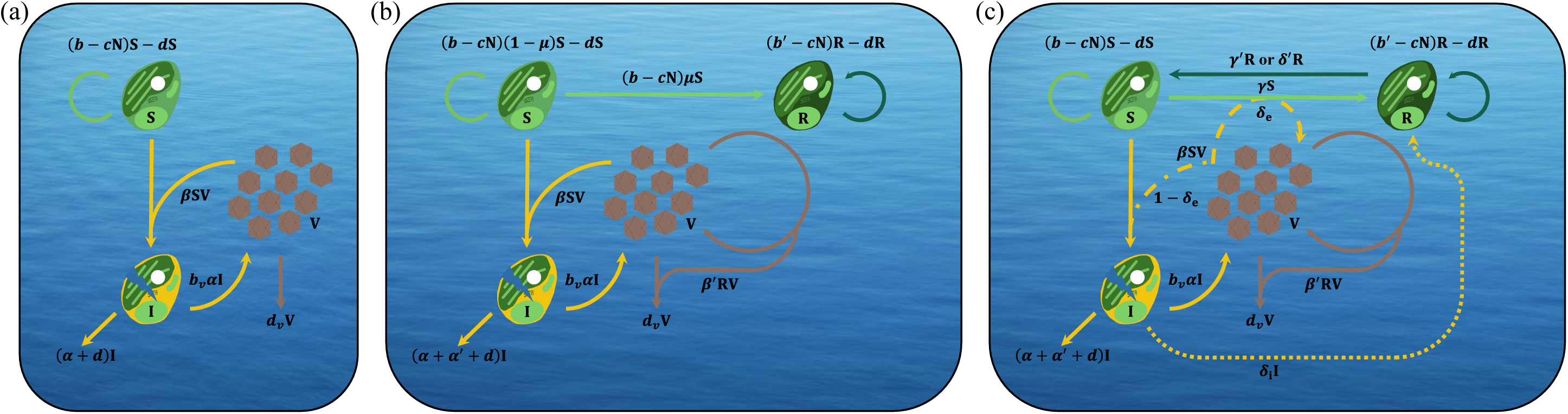
Modelling of the phytoplankton-virus interaction dynamics. (a) SIV model without antiviral resistance. (b) SRIV-GM model with antiviral resistance associated to Genomic Mutations. (c) SRIV-RPP and SRIV-IPP models with antiviral resistance acquired by Random or virus-Induced Phenotypic Plasticity. S, R, I stand for the density of susceptible, resistant, and infected host cells, while V denotes the density of virions in the marine environment. All parameters are defined in the main text, and their estimates are provided in table 1. In (c) continuous, dashed, and dotted lines describe the processes specific to random, extracellular induced and intracellular induced phenotypic plasticity, respectively. The specific representations of Fig. 1c corresponding to SRIV-RPP, SRIV-IPP with extracellular and intracellular resistance mechanisms are provided in Supporting Information Fig. S1.

**Table 1.**
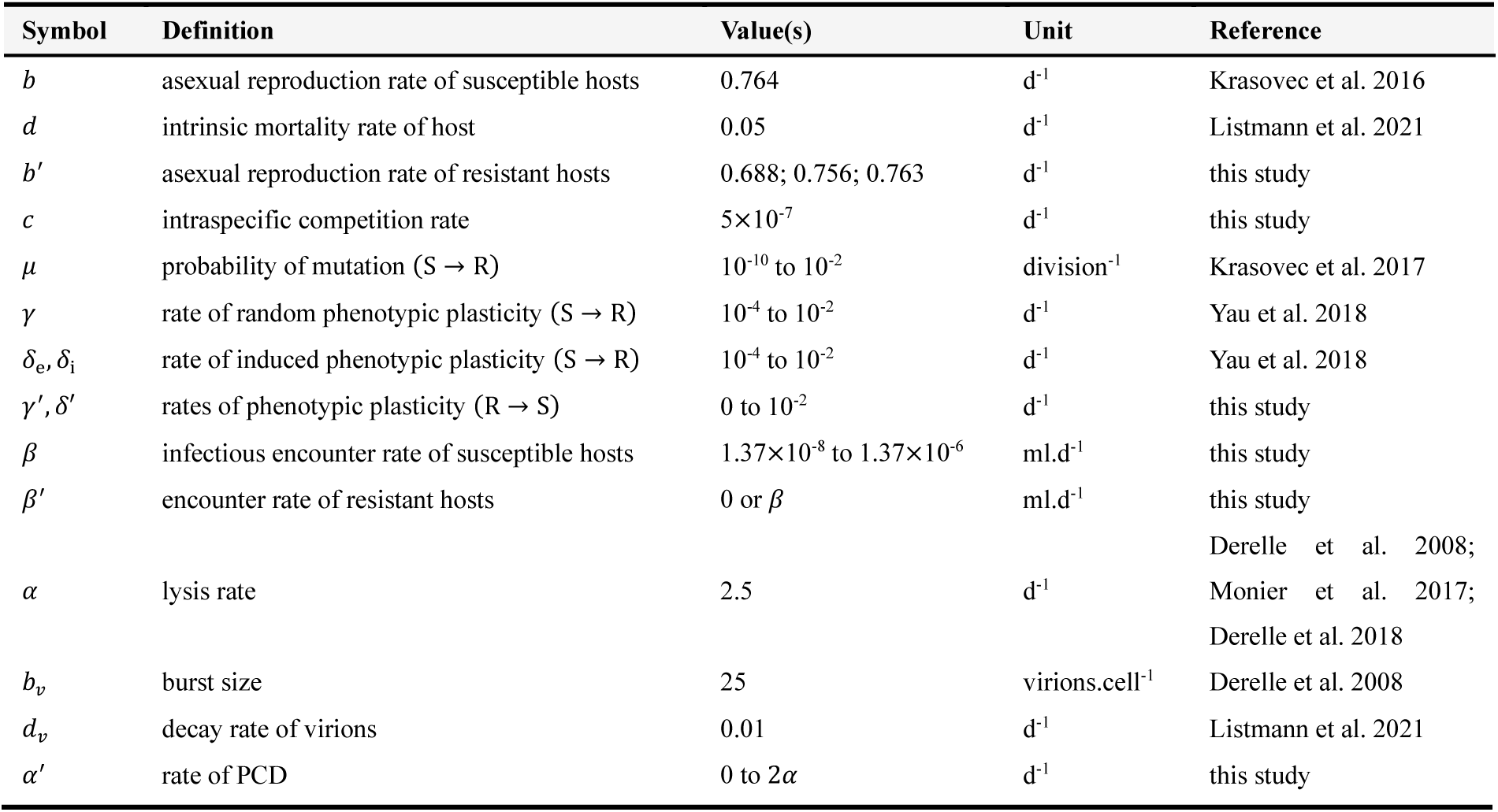
Estimates of the SIV-SRIV modelling parameters describing O. tauri-OtV interactions. The definition of all modelling parameters describing O. tauri and OtV life-history traits, their encounter rates, as well as the determinism and mechanisms of antiviral resistance are provided with the corresponding symbols, values, and units. A detailed explanation of the estimates is provided in Supporting Information, Heading 1.

| Symbol | Definition | Value(s) | Unit | Reference |
|---|---|---|---|---|
| *b* | asexual reproduction rate of susceptible hosts | 0.764 | d<sup>-1</sup> | Krasovec et al. 2016 |
| *d* | intrinsic mortality rate of host | 0.05 | d<sup>-1</sup> | Listmann et al. 2021 |
| *b*′ | asexual reproduction rate of resistant hosts | 0.688; 0.756; 0.763 | d<sup>-1</sup> | this study |
| *c* | intraspecific competition rate | 5×10<sup>-7</sup> | d<sup>-1</sup> | this study |
| *μ* | probability of mutation (S → R) | 10<sup>-10</sup> to 10<sup>-2</sup> | division<sup>-1</sup> | Krasovec et al. 2017 |
| *γ* | rate of random phenotypic plasticity (S → R) | 10<sup>-4</sup> to 10<sup>-2</sup> | d<sup>-1</sup> | Yau et al. 2018 |
| *δ*<sub>e</sub>, *δ*<sub>i</sub> | rate of induced phenotypic plasticity (S → R) | 10<sup>-4</sup> to 10<sup>-2</sup> | d<sup>-1</sup> | Yau et al. 2018 |
| *γ*′, *δ*′ | rates of phenotypic plasticity (R → S) | 0 to 10<sup>-2</sup> | d<sup>-1</sup> | this study |
| *β* | infectious encounter rate of susceptible hosts | 1.37×10<sup>-8</sup> to 1.37×10<sup>-6</sup> | ml.d<sup>-1</sup> | this study |
| *β*′ | encounter rate of resistant hosts | 0 or *β* | ml.d<sup>-1</sup> | this study |
| *α* | lysis rate | 2.5 | d<sup>-1</sup> | Derelle et al. 2008; Monier et al. 2017; Derelle et al. 2018 |
| *b*<sub>*v*</sub> | burst size | 25 | virions.cell<sup>-1</sup> | Derelle et al. 2008 |
| *d*<sub>*v*</sub> | decay rate of virions | 0.01 | d<sup>-1</sup> | Listmann et al. 2021 |
| *α*′ | rate of PCD | 0 to 2*α* | d<sup>-1</sup> | this study |

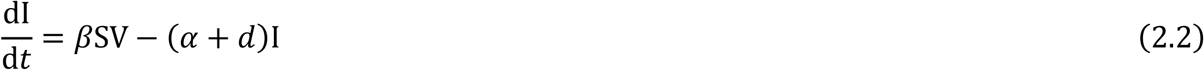

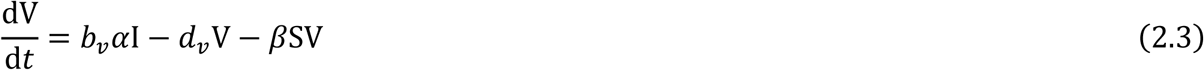

that served as a core model to further account for antiviral resistance and its alternative determinisms and mechanisms. The definition of all parameters of this SIV core model and of its SRIV extensions, as well their estimates for *O. tauri* and *O. tauri* viruses (OtV), are provided in the summary table 1.

### **SRIV** modelling of phytoplankton’s antiviral resistance

To investigate the role of antiviral resistance on the population dynamic of phytoplankton-virus interaction, we integrated a compartment of resistant hosts into our core model. The production of such individuals obviously depends on the determinism of resistance, which led to design two separate models under the genomic mutations (Fig. 1b) and (random or virus-induced) phenotypic plasticity (Fig. 1c) hypotheses. In both of these SRIV models, where R denotes the density of resistant cells, the mechanisms of (extracellular or intracellular) individual resistance and PCD were described as changes in the rates of encounter or lysis.

### **Genomic** mutations for antiviral resistance (GM)

When resistance is acquired through genomic mutations (Fig. 1b), the asexual division of susceptible cells can produce either resistant cells, according to the mutation probability *μ*, or susceptible cells, with probability (1 - *μ*). Resistant cells then reproduce with respect to specific rates *b*’ = *b*(1 – ζ), where ζ stands for the individual cost of resistance. All daughter cells are then considered to be resistant ones as reverse mutations toward susceptibility are highly unlikely.

The outcome of encounters between resistant cells and virions was described according to the mechanism of antiviral resistance considered. In the case of an extracellular resistance, the virus typically does not adsorb onto the host cell and merely returns to the environmental pool of virions, which effectively makes the encounter rate between a resistant cell and a virion (*β*^′^) equal to 0. On the contrary, when antiviral resistance is assumed to be intracellular, the encounter rate does not differ from that of a susceptible cell (*β*^′^ = *β*). Since the virus introduces its genetic material into the resistant cell, which is then able to control its within-host dynamic, such contacts effectively lead to remove virions from the environmental pool of infective viral particles at rate *β*^′^RV.

In addition to the above mechanisms of individual resistance, infected cells can also trigger PCD, which is expected to provide a collective defence by limiting the production of virions. Such a mechanism is therefore not associated with resistant cells and instead can be described by an additional mortality rate (*α*’) of infected cells allowing to avoid viral lysis and the associated release of virions.

The descriptions of phytoplankton-virus interactions with antiviral resistance acquired through genomic mutations (Fig. 1b) therefore led to the following SRIV-GM model;

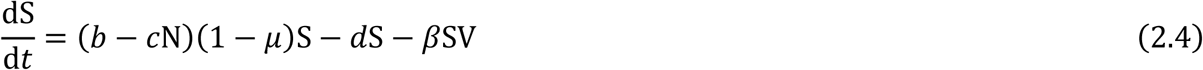

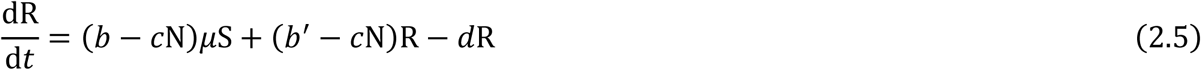

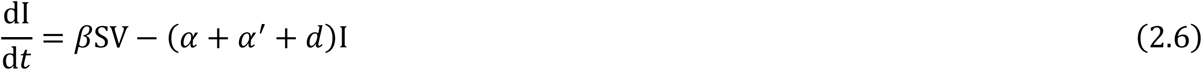

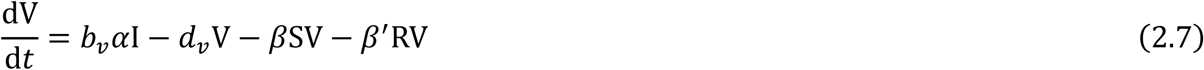

where the size of the phytoplankton population N = S+R+I.

### **Phenotypic** plasticity between susceptibility and antiviral resistance (PP)

When resistance is acquired through phenotypic plasticity (Fig. 1c), the cells’ switches from a susceptible to a resistant phenotype can occur randomly (RPP, Fig. 1c) as a result of a ‘bet-hedging’ strategy, or be induced by viruses (IPP, Fig. 1c).

Under the RPP hypothesis, susceptible cells can become resistant at a constant rate (*γ*) and resistant cells can switch back to a susceptible phenotype at a similarly constant rate (*γ*’). Since such phenotypic changes happen throughout the cell lifetime, they are not directly associated with asexual reproduction. The outcome of encounters between resistant cells and virions was then modelled according to the (extracellular and intracellular) mechanisms of antiviral resistance, considered exactly as under the hypothesis that antiviral resistance arises from genomic mutations. The inclusion of PCD also remained unchanged and accounted for as an additional mortality rate of infected cells (*α*’). The introduction of a random phenotypic plasticity between susceptibility and resistance into our core model (Fig. 1c) therefore led to the following SRIV-RPP model;

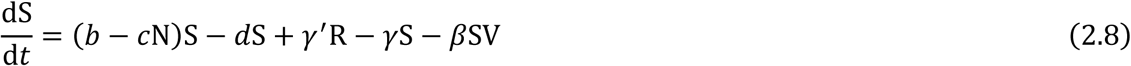

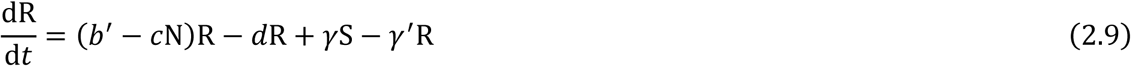

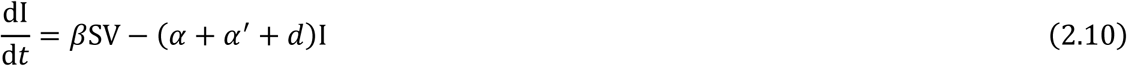

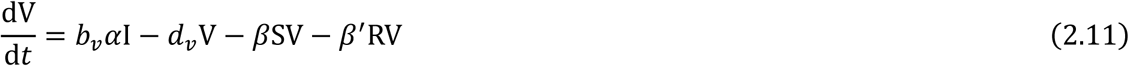

Under the IPP hypothesis, the timing of the induction of antiviral resistance changes with respect to the mechanism considered. When the resistance is extracellular, the contact between a susceptible cell and a virus triggers the phenotypic change. A proportion (*δ*_e_) of such contacts was then assumed to produce resistant cells while others lead to infection. As previously explained, no viral mortality was associated with such an extracellular resistance (*β’ =* 0) since virions merely stay in the environmental pool. When the resistance is intracellular, the virion still adsorbs onto the susceptible host and the phenotypic switch to resistance is induced through the infection. Infected cells were then considered to become resistant at a constant rate (*δ*_i_). As described earlier, further contacts between intracellular resistant cells and virions led to the removal of the latter from the environmental pool (at rate *β’* = *β*). In both the extracellular and intracellular cases, resistant cells can switch back to a susceptible phenotype at a similarly constant rate (*δ*^′^). These dynamics of phytoplankton-virus interactions with phenotypic switches induced by infection (Fig. 1c) can then be predicted by the following SRIV-IPP model;

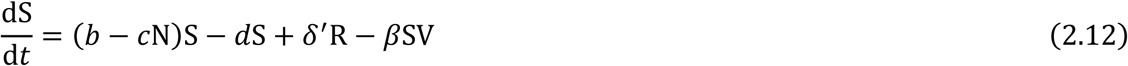

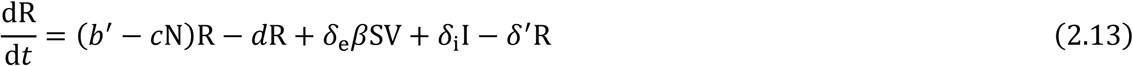

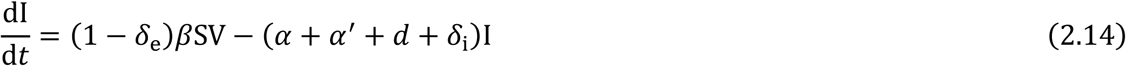

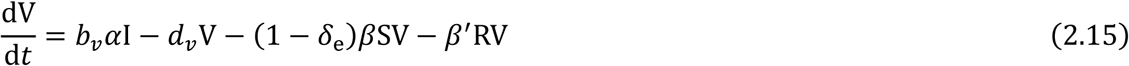

The two specific forms of this model that we used to investigate the impacts of either extracellular (*δ*_e_, *δ*_i_=0) or intracellular (*δ*_e_=0, *δ*_i_) resistance on the phytoplankton-virus interaction dynamics are provided in Supporting Information Fig. S1.

### **Theoretical** analyses of the impact of antiviral resistance strategies

We investigated the impacts of antiviral resistance on the population dynamics of phytoplankton-virus interactions through standard local stability analyses of each of our SRIV-GM, SRIV-RPP and SRIV-IPP models. Such analyses performed on epidemiological models typically led to the identification of three equilibrium points informing on the densities of host and virus individuals that can be expected under different conditions; a Trivial Equilibrium (TE) where neither (phytoplankton) hosts nor viruses are present, a Disease-Free Equilibrium (DFE), and a Disease Equilibrium (DE), where the hosts is present without or with the viruses. Those analyses further allowed to identify i) the so-called 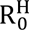 of the host measuring the rate at which the phytoplankton is able to grow when it develops in a virus-free environment, ii) the 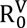 of its virus, providing the rate at which it can spread in a phytoplankton host population at its DFE, and iii) when the host-virus interactions lead to their coexistence at a stable DE or through oscillatory dynamics around this equilibrium. We used those general methods to answer the three key questions raised in the introduction about the effects of antiviral resistance on the main stages of phytoplankton-virus ecological dynamics, as described in more details below.

### **How** does (the cost of) antiviral resistance affect phytoplankton’s population?

We first considered a virus-free environment in order to predict the effect of the cost of resistance on the intrinsic growth rate of a phytoplankton population (R^H^) and on its potential biotic capacity, i.e. its equilibrium level at the DFE (K_N_). Such effects were measured as a percentage of reduction in 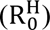 and K_N_ resulting from the introduction of a cost of resistance reducing the reproductive rate of resistant cells. Those were assessed for different costs (ζ = 0.1%, 1%, 10%) and, according to the antiviral resistance determinism, for ranges of mutation probability (*μ* in [10^-10^;10^-2^]) and rates of random phenotypic plasticity (*γ* in [10^-4^;10^-2^] and *γ*’ in [0;10^-2^]). To do so, for each of our SRIV models, we derived the expression of the phytoplankton 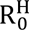 (by identifying the conditions for the phytoplankton population to grow, i.e. when the TE is unstable), and its potential biotic capacity (by determining the expected abundance of phytoplankton at the DFE).

### **How** does the resistance limit the initial spread of the virus?

We then looked at the impact of antiviral resistance on the rate of spread of a virus emerging in a phytoplankton population that has reached its potential biotic capacity (K_N_). Such an impact was defined as the percentage of reduction in the 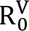 of the virus that is induced by the existence of antiviral resistance. This was assessed for the same costs of resistance (ζ), mutation probabilities (*μ*) and rates of random phenotypic plasticity (*γ* and *γ*’) as described in section 2.2.1, and by varying the rates of induced phenotypic plasticity (*δ*_e_ and *δ*_i_ in [10^-4^;10^-2^] and *δ’* in [0;10^-2^]), while considering several rates of PCD (*α*’ in [0,5]). For each of our SRIV models, the expression of 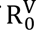 was derived as the conditions allowing for the virus to spread in a phytoplankton population at its biotic capacity (by determining when the DFE is unstable).

### **How** does resistance influence the stability of the host-virus interaction and the ‘top-down’ control of the phytoplankton?

We investigated the potential contribution of each determinism and mechanism of antiviral resistance to the dynamics of the phytoplankton-virus interaction. To determine when they coexist at a stable DE or by oscillating around this equilibrium, we looked at the stability of the DE using the Routh-Hurwitz criteria (Otto and Day 2011) and by assessing them for costs of resistance (ζ) varying from 0 to 10%. We used the same ranges of mutation probability (*μ*), random (*γ* and *γ*’) or induced (*δ*_e_ and *δ*_i_) rates of phenotypic plasticity, and rates of PCD (*α*’) as described in the above sections.

### **Tailoring** of the model parameters to *Mamiellophyceae* – prasinoviruses interactions

The stability analyses of our SRIV models allowed to identify the expressions of their (DFE and DE) equilibrium and the R_0_ of both hosts and viruses. While these general expressions can allow investigating the impact of antiviral resistance on the dynamics of interactions in various biological systems made of a unicellular host and its lytic virus, we used them to make quantitative predictions about the effect of antiviral resistance on three main stages of the phytoplankton-virus population dynamics. We then focused our investigations on a key *Mamiellophyceae* species, *O. tauri*, that has been repeatedly shown to exhibit antiviral resistance to its highly specific lytic prasinoviruses, OtV (Thomas et al. 2011; Yau et al. 2016, 2018; Thomy et al. 2026). Previous experimental studies of *O. tauri-*OtV interactions allowed to derive estimates of both the phytoplankton and viruses’ life-history traits, and to specify the relevant ranges for the encounter rate as well as the parameters describing antiviral resistance and collective defence in each of our SRIV models. Those estimates and ranges of parameter values used in all numerical analyses are summarised in table 1.

## Results

### **How** does antiviral resistance affect phytoplankton’s population?

We first investigated the effects of the cost of resistance on the intrinsic growth rate of a phytoplankton population 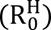 and on its potential biotic capacity (K_N_) in a virus free environment. The expressions of the phytoplankton’s i) 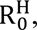 ii) population size K_N_ and iii) amount of resistant cells (R*) reached at the DFE, were then derived under the different hypotheses about the determinism and mechanism of antiviral resistance (Supporting Information, Headers 2-4).

As expected, none of these quantities were found to depend on (the parameters *β*^′^ and *α*’ describing) the mechanisms of resistance and PCD since, in the absence of virus, neither individual resistance nor a collective defence can be triggered. The only predicted differences occur according to the determinism of resistance (table 2). Under the hypothesis of an induced phenotypic plasticity (IPP), the lack of virus in the environment obviously results in the absence of resistant cells. The phytoplankton’s 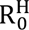 and K_N_ were then found to be the same as those derived for the SIV core model, which led to an absence of effect on the phytoplankton population. Under the GM and RPP hypotheses, the spontaneous production of antiviral resistant cells reduces both 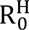 and K_N_, albeit to a different extant.

**Table 2.** Expressions of the phytoplankton’s intrinsic growth rate 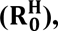 potential biotic capacity (KN) and the proportion of resistant cells (R*/KN) at the disease-free equilibrium. Expressions are provided for the three potential determinisms of antiviral resistance; Genomic Mutations (GM), random phenotypic plasticity (RPP), and virus-induced phenotypic plasticity (IPP). These were found to be independent of the (extracellular .vs. intracellular) mechanisms of resistance.

| **Genomic Mutations (GM)** | **Random Phenotypic Plasticity (RPP)** | **Induced Phenotypic Plasticity (IPP)** |
|---|---|---|
| $\mathrm{R}_0^{\mathrm{H}} = \frac{b(1-\mu)+b'}{2d}$ | $\mathrm{R}_0^{\mathrm{H}} = \frac{b+b'}{2d+\gamma+\gamma'}$ | $\mathrm{R}_0^{\mathrm{H}} = \frac{b}{d}$ |
| $\mathrm{K}_\mathrm{N} = \frac{1}{c}\left(b - \frac{d}{(1-\mu)}\right)$ | $\mathrm{K}_\mathrm{N} = \frac{r + r' - \gamma - \gamma' + \sqrt{b\zeta^2 + (\gamma+\gamma')^2 + 2b\zeta(\gamma'-\gamma)}}{2c}$ | $\mathrm{K}_\mathrm{N} = \frac{r}{c}$ |
| $\frac{\mathrm{R}^*}{\mathrm{K}_\mathrm{N}} = \frac{d\mu}{b\zeta(1-\mu)}$ | $\frac{\mathrm{R}^*}{\mathrm{K}_\mathrm{N}} = \frac{\gamma}{c\mathrm{K}_\mathrm{N} + \gamma + \gamma' - r'}$ | $\frac{\mathrm{R}^*}{\mathrm{K}_\mathrm{N}} = 0$ |

We then used these general metrics to quantify the effects of antiviral resistance on a *O. tauri* population growing in a virus free environment, by estimating the reduction of its intrinsic growth rate 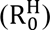 and expected biotic capacity (K_N_) under the GM and RPP hypotheses (Fig. 2).

**Figure 2.**
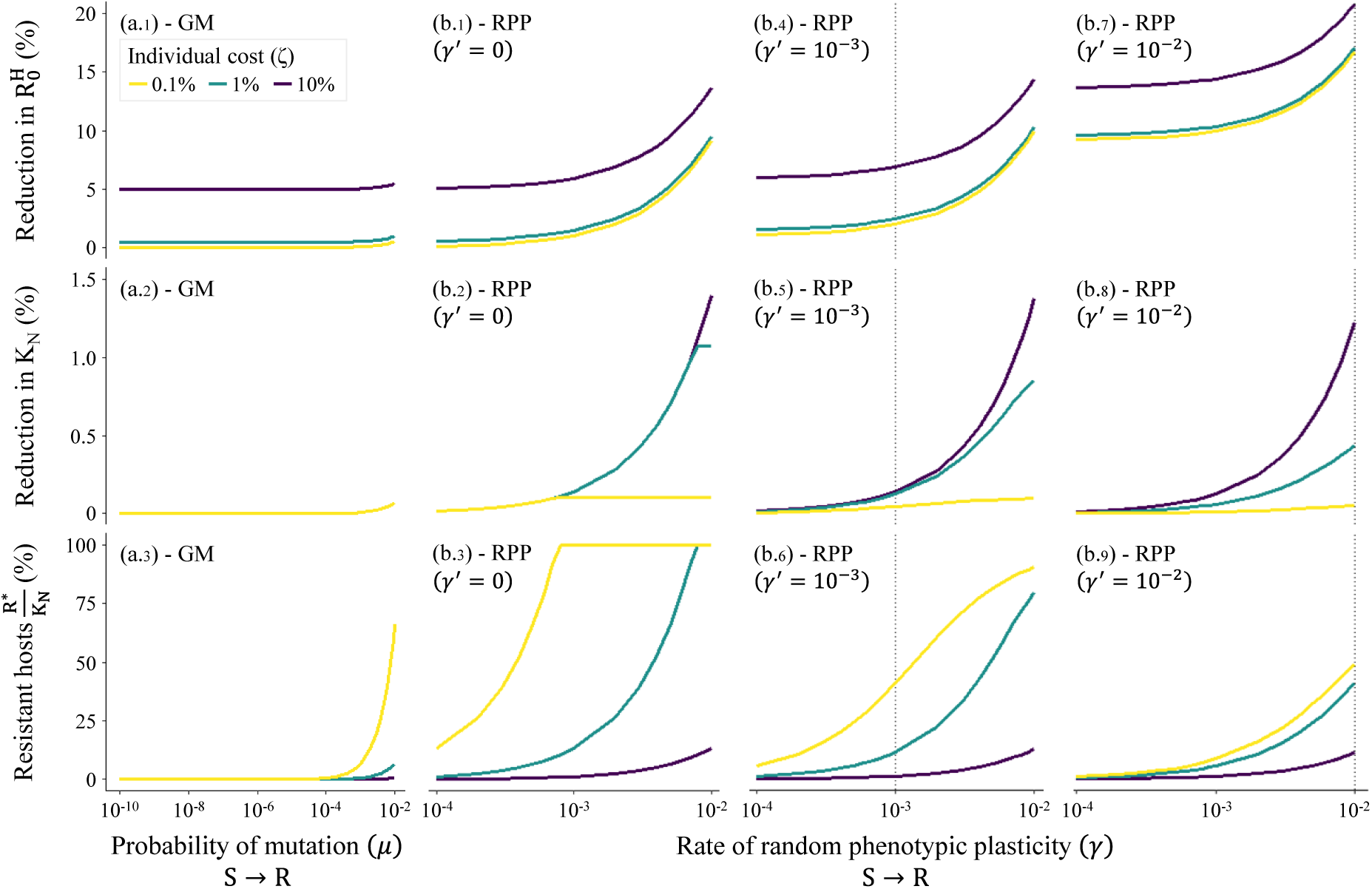
Effects of antiviral resistance on the *O. tauri*’s intrinsic growth rate 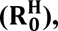 biotic capacity (K_N_) and percentage of resistant cells (R*/K_N_). (a) Antiviral resistance associated to Genomic Mutations (SRIV-GM model). (b) Antiviral resistance acquired through Random Phenotypic Plasticity (SRIV-RPP model). The effects on 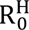 and K_N_, along with those on the percentage of resistant cells in the population of *O. tauri*, are shown according to the probability of mutation (*μ*) or the rates of random phenotypic plasticity (*γ*) toward antiviral resistance when such resistance is determined by GM (a.1-3) and RPP (b.1-9), respectively. Under the RPP hypothesis, the rate at which resistant can switch back to a susceptible phenotype (*γ’*) is set to 0 (b.1*-*3), 10^-3^ (b.4*-*6) and 10^-2^ (b.7*-*9). In each panel, the impact of antiviral resistance is shown according to an individual cost of resistance (*ζ*) equals to 0.1% (yellow), 1% (green) and 10% (purple).

When antiviral resistance occurs as a result of genomic mutations (GM), its effect on the intrinsic growth rate of the phytoplankton (R^H^) remains typically lower than 1%, unless its individual cost (*ζ*) takes on a very high value that led to population effects reaching up to 5.5% (Fig. 2a.1). Its impact on the maximal abundance potentially reached by *O. tauri* population is even smaller as it always remains below 0.1% when the probability of mutation is set to be very high (10^-2^), whatever the individual cost of resistance (Fig. 2a.2). The percentage of resistant cells then typically remains below ∼7%, unless the cost of resistance is very low (Fig. 2a.3).

The existence of a random phenotypic plasticity (RPP) providing resistant cells led to larger variations of the *O. tauri* intrinsic growth rate and biotic capacity (Fig. 2b.1-9). The reduction of the intrinsic growth rate (R^H^) then rises with the rate at which resistance is acquired to reach up to ∼9 to 14%, depending on the individual cost of resistance (Fig. 2b.1). Although such reduction is ∼3 to 17 times larger than when antiviral resistance is associated with a genomic mutation, the impact on *O. tauri*’s biotic capacity (K_N_) always remains lower than 1.5% (Fig. 2b.2). Such a maximal reduction in *O. tauri*’s K_N_ is obviously reached when the individual cost of resistance is high and, accordingly, the percentage of resistant cells is low (Fig. 2b.3). When the cost of resistant is reduced, this percentage can increase very significantly, although the impact on the phytoplankton abundance progressively vanishes (Fig. 2b.2-3). As expected from the general expressions of 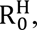 increasing the rate of switch reversing resistant into susceptible cells (*γ*’) enhances the impact of antiviral resistance on the phytoplankton’s 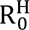 by reducing the expected amount of time available for asexual reproduction over the cell’s lifetime, i.e 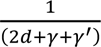 (Fig. 2b.4,b.7). Meanwhile, such an increase reduces the effect of resistance on the biotic capacity (K_N_) of *O. tauri* (Fig. 2b.5,b.8) since it obviously lowers the percentage of resistant cells in the population (Fig. 2b.6,b.9).

### **How** does the resistance limit the initial spread of the virus?

We then looked at the effects of antiviral resistance focusing on how efficient it can be to reduce the initial rate of spread of a virus 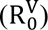 emerging in a phytoplankton population that has reached its potential biotic capacity (K_N_), i.e. at the DFE.

Noteworthy, the expressions of the 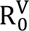 identified under each of our hypotheses about the determinism and mechanism of antiviral resistance (Supporting Information, Headers 2-4) are all defined by three terms that neatly unravel how those hypotheses impact the production of ‘secondary’ virions through their effects on each key stage of the viral cycle (table 3). The first term gives the rate at which a ‘primary’ virion infects a susceptible host, which typically depends on the density of susceptible individuals (S*) and the intrinsic rate of encounter (*β*). The second corresponds to the net number of virions released from a ‘primary’ infection, which is defined with respect to both the probability for the infected host to be lysed (*α*) before it dies of natural causes (*d*) or PCD (*α*’), and the number of virions subsequently released in the environment (*b_v_*). Finally, since the above rate of ‘primary’ infection is defined per unit time, it has to be weighted by the expected lifetime of the ‘primary’ virion in the environment, which is inversely related to the sum of its rate of mortality by natural causes (*d_v_*) and encounters with intracellular resistant cells present at the DFE (*β*^′^R*). Altogether, those three terms provide meaningful expressions of the expected production of ‘secondary’ virions from a single ‘primary’ one, i.e. of the virus basic reproduction number 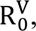, that clearly depend on both the determinism and the mechanism of antiviral resistance.

**Table 3.** Expressions of the virus basic reproduction number 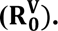. The expression of 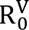 is similar when antiviral resistance is associated with Genomic Mutations (GM) and random phenotypic plasticity (RPP), while it differs when it is acquired through virus-induced phenotypic plasticity (IPP) and corresponds to an extracellular or an intracellular mechanism.

| **Genomic Mutations and Random Phenotypic Plasticity** | **Induced Phenotypic Plasticity (Ext.)** | **Induced Phenotypic Plasticity (Int.)** |
|---|---|---|
| $\mathrm{R}_0^\mathrm{V} = \beta \mathrm{S}^* \left(\frac{\alpha}{(\alpha + \alpha' + d)} b_v - 1\right) \frac{1}{d_v + \beta' \mathrm{R}^*}$ | $\mathrm{R}_0^\mathrm{V} = (1 - \delta_\mathrm{e}) \beta \mathrm{S}^* \left(\frac{\alpha}{(\alpha + \alpha' + d)} b_v - 1\right) \frac{1}{d_v}$ | $\mathrm{R}_0^\mathrm{V} = \beta \mathrm{S}^* \left(\frac{\alpha}{(\alpha + \alpha' + d + \delta_\mathrm{i})} b_v - 1\right) \frac{1}{d_v}$ |

We then used these general expressions of 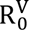 to look at the expected effects of antiviral resistance on the initial growth rate of OtV emerging in a *O. tauri* population that has reached the DFE under the GM, RPP and IPP hypotheses (Fig. 3).

**Figure 3.**
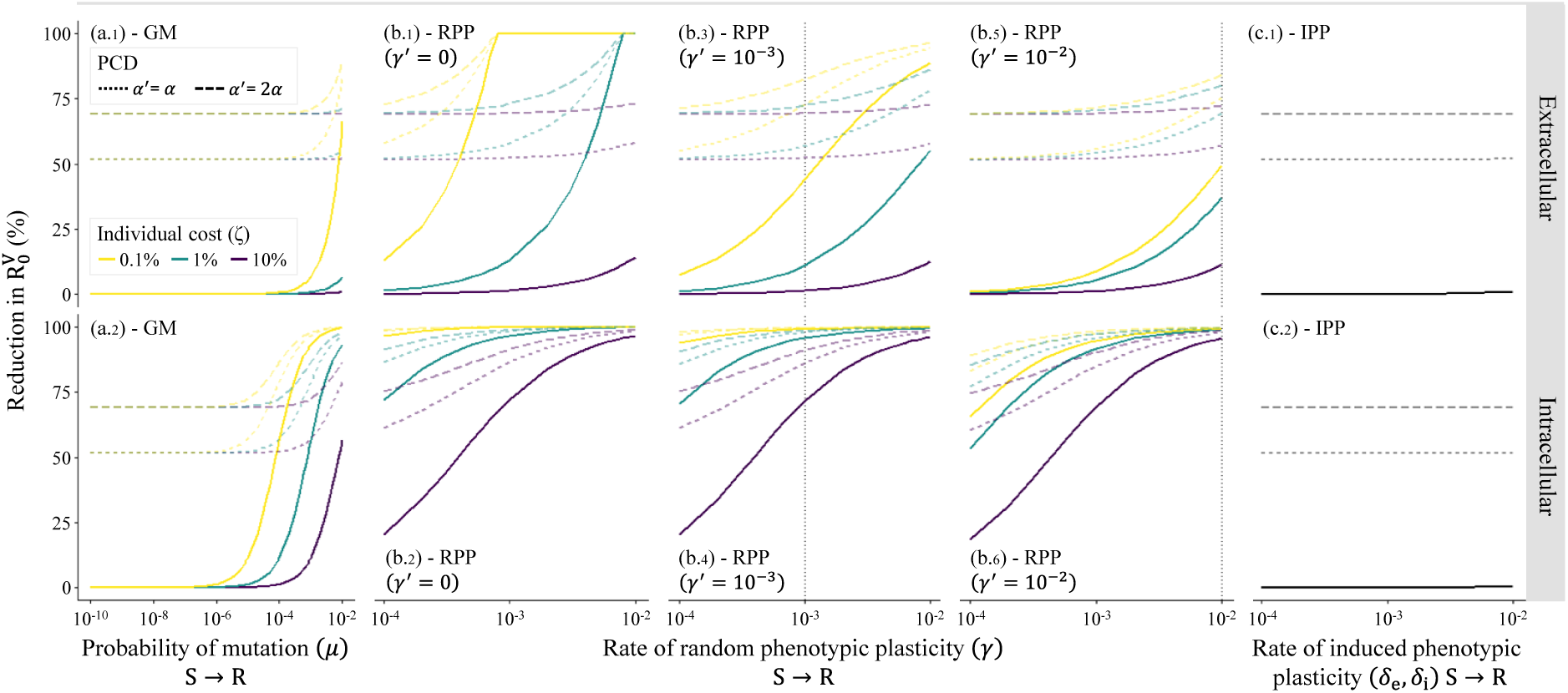
Effects of antiviral resistance on the basic reproductive number of OtV 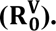. (a) Antiviral resistance associated to Genomic Mutations (SRIV-GM model). (b) Antiviral resistance acquired through Random Phenotypic Plasticity (SRIV-RPP model), with a rate of phenotypic switch from R to S (*γ*^′^) equals to zero (b.1-2), 10^-3^ (b.3-4) or 10^-2^ (b.5-6). (c) Antiviral resistance acquired through virus Induced Phenotypic Plasticity (SRIV-IPP model).

For a, b and c, the effects of antiviral resistance are shown when it corresponds to an extracellular (first row) or an intracellular (second row) mechanism, and in the absence (solid lines) or presence of PCD (dashed lines: *α*′=*α*, long dashed lines: *α*′=2*α*). In each of the a and b panels, yellow, green and purple lines show the effects of antiviral resistance when its individual cost (*ζ*) is equal to 0.1%, 1% and 10%, respectively. In panels c.1 and c.2, such a cost has no impact since there are no resistant cells in the initial stage of the viral spread.

Antiviral resistance associated with genomic mutations (GM) and extracellular mechanisms shows a limiting effect on 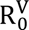 (Fig. 3a.1 with no PCD, solid lines). This effect tightly varies with the proportion of resistant cells at the DFE (see Fig. 2a.3), since increasing R* directly reduces the density of susceptible cells available for infectious contact (*β*S*). Such a ‘dilution’ effect can only limit the spread of OtV when the mutation probability is high (*μ* > ∼10^-4^). On the contrary, the collective defence provided by PCD appears much more efficient as it allows halving 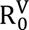 (Fig. 3a.1, dashed lines) as soon as it induces the death of infected cells as fast as lysis (i.e. for *α*’=*α* and 2*α*). As expected, such an effect of PCD manifests itself even at low mutation probabilities as it is not directly related to resistant cells. Interestingly, the effect of antiviral resistance on the 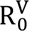 is substantially higher when such resistance is intracellular (Fig. 3a.2) since virions are then removed from the environmental pool of infective particles. The emergence of OtV then starts to be limited by *O. tauri* resistance for much lower mutation probabilities, i.e. for *μ* between 10^-7^ to 10^-5^, depending on the individual cost of resistance. As expected, the efficacy of the PCD was the same as for extracellular case, as it is independent of the resistance mechanism at work.

When random phenotypic plasticity (RPP) allows for the production of resistant cells through an extracellular mechanism that prevents viral adsorption (Fig. 3b.1 with no PCD, solid lines), the ‘dilution’ effect still matches the variation in R* at the DFE (see Fig. 2b.3). Since the rate of phenotypic switch to resistance (*γ*) is typically higher than the mutation probability from S to R (*µ*), the predicted effects of RPP on the spread of OtV are understandably larger. However, the existence of reverse phenotypic switches towards susceptibility (Fig. 3b.3,b.5) can substantially reduce the impact of antiviral resistance on the spread of OtV to almost similar levels as those predicted under the GM hypothesis. As explained above, the collective defence provided by PCD does not depend on the strategy of individual resistance and the rate at which it appears (*γ*). Accordingly, its effect (Fig. 3b.1, dashed lines) is again dominant when *γ* is low and can at least halve the 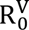 (as soon as *α*’=*α*). Meanwhile, intracellular resistance is still much more efficient in limiting the spread of OtV because virions entering the resistant cells are wasted (Fig. 3b.2,b.4,b.6) so that the reduction of the 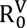 is predicted to be almost always larger than 50%.

The effect of antiviral resistance induced by the virus (IPP) on its own rate of emergence is typically low for both extracellular (Fig. 3c.1) and intracellular (Fig. 3c.2), provided that 1 − *δ*_e_ ≃ 1 and *α* + *α*^′^ + *d* + *δ*_i_ ≃ *α* + *α*^′^ + *d* (since *δ*_e_, *δ*_i_ ≤ 10 − 2) as it could be anticipated from the general expressions of 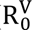 appearing in table 3.

### **How** does resistance influence the stability of the host-virus interaction and the ‘top-down’ control of the phytoplankton?

While it is well known that phytoplankton and virus coexist in marine environments, how antiviral resistance influences the dynamical outcome of their interaction remains unclear. Using the (general) outcomes of the stability analyses of the DE that we identified for each of our SRIV models (Supporting Information, Headers 2-4), we predicted when coexistence between *O. tauri* and OtV is expected to occur through typical host-virus oscillations and when individual resistance and PCD can promote a stable coexistence at such DE.

Noteworthily, the dynamical regime of *O. tauri*-OtV coexistence was only weakly affected by the (extracellular .*vs*. intracellular) mechanism of resistance. We then focus (below and in Fig. 4) on the intracellular case, and we refer to Supporting Information Fig. S2 for a similar description of the impacts of extracellular antiviral resistance.

**Figure 4.**
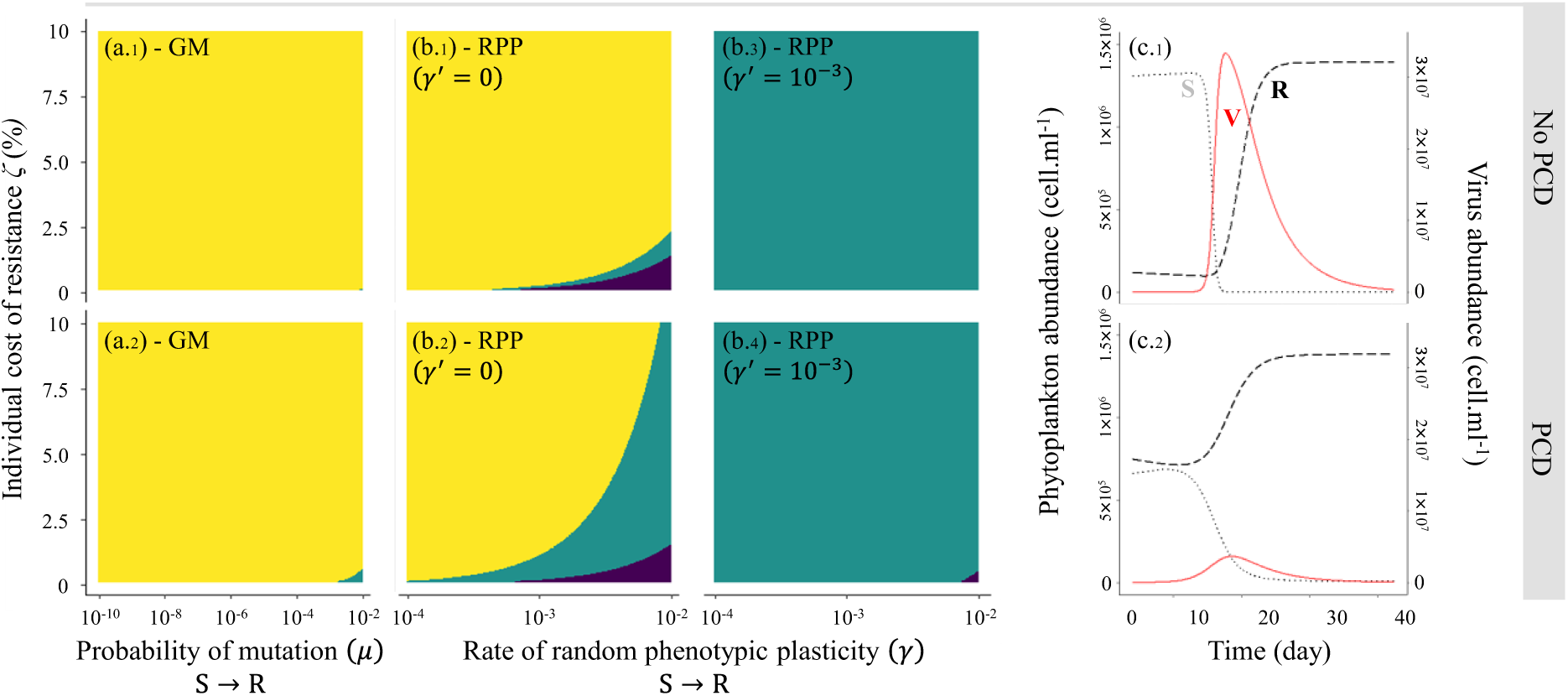
Effects of antiviral resistance on the dynamics of *O. tauri*–OtV interactions. (a) Antiviral resistance associated to Genomic Mutations (SRIV-GM model). (b) Antiviral resistance acquired through Random Phenotypic Plasticity (SRIV-RPP model) with a rate of phenotypic switch from R to S (*γ*^′^) equal to zero (b.1-2) and 10^-3^ (b.3-4). (c) Variations of S-R host cells and virions V abundances during the ‘epidemic’ stage of a typical host-virus oscillatory dynamics. For a, b and c, the outcomes of the *O. tauri*-OtV interaction are predicted in the absence (first row) or in the presence (second row) of PCD (with *α*′ =2*α*). In each a and b panels, yellow and green areas correspond to *O. tauri*–OtV coexistence through oscillatory and stable dynamics, respectively, while purple areas show the conditions where the OtV is unable to spread.

The dynamical outcome of *O. tauri*-OtV interactions predicted when resistance is associated with genomic mutations (GM) corresponds to the so-called ‘boom and bust’ dynamics (Flynn et al. 2022) for the broad range of probability of mutation (*μ*) and individual cost of resistance (*ζ*) considered (Fig. 4a.1). This oscillatory dynamic is typically made of an ‘epidemic’ phase (Fig. 4c.1) starting when the susceptible *O. tauri* population has recovered from the previous virus-induced ‘bust’. The number of infections and virions then rises sharply until the susceptible *O. tauri* population is nearly depleted and the remaining population is dominated by resistant individuals. This is followed by a basic ‘recovery’ phase (not shown) whereby a reduced density of OtV virions leads to a ‘boom’ of the susceptible part of the *O. tauri* population, which allows initiating the next ‘epidemic’ phase. Increasing the probability of mutation (*μ*) or lowering its individual cost (*ζ*), i.e. going towards the bottom right corner of Fig. 4a.1, obviously enhances the density of resistant *O. tauri* cells during the ‘epidemic’ phase. This lowers the peak of OtV abundance and dampens the oscillations but, in the absence of PCD, does not allow for a stable equilibrium to be reached (Fig. 4a.1). As expected, the existence of a PCD significantly reduces the number of infections and virions during the ‘epidemic’ phase (Fig. 4c.2). Interestingly, this allows the susceptible *O. tauri* cells, although still strongly reduced, to persist at higher abundances at the end of this ‘epidemic’ stage and to sustain a larger pool of OtV, which ultimately limits the re-building of this susceptible part of the host population. This explains why, at the start of the ‘epidemic’ phase, less susceptible cells are available, which further contributes to reduce the number of infections and to dampen the *O. tauri*-OtV oscillations. Despite these direct and indirect effects of PCD, it still did not allow stabilising the interaction dynamics, unless a very high probability of mutation and a low individual cost of resistance allowed for resistant cells to remain consistently dominant in the population (Fig. 4a.2).

When individual resistance is acquired by random phenotypic plasticity (RPP) and resistant cells cannot switch back to a susceptible phenotype (*γ*’ = 0), *O. tauri*-OtV oscillations still prevail for a broad range of random rates of phenotypic switch (*γ*) and individual cost of resistance (*ζ*), although a stable coexistence can be reached under a substantial range of conditions (Fig. 4b.1,b.2). The dynamics of oscillations indeed tend to be of lower amplitudes, as compared to those predicted with the GM model, since the typically higher rate of phenotypic plasticity allows for resistant *O. tauri* cells to be produced faster during the build-up of the peak of OtV characterising the ‘epidemic’ phase. In the absence of PCD (Fig. 4b.1), when the individual cost of resistance is lower than 1.5%, high rates of phenotypic switch (*γ*) can turn the oscillatory dynamics of *O. tauri*-OtV interaction (yellow) into a stable coexistence (green) and even prevent the spread of OtV (purple). Interestingly, the co-occurrence of an antiviral resistance acquired through RPP and PCD facilitates *O. tauri*-OtV stable coexistence, which become feasible for rates of phenotypic switch *γ* < 10^-3^ or individual cost to resistance *ζ* > 1% (Fig. 4b.2). Strikingly, when resistant cells can switch back to a susceptible phenotype (*γ*’ > 0), the *O. tauri*-OtV interactions are almost always predicted to lead to a stable coexistence (Fig. 4b.3,b.4). The switch back from resistant to susceptible phenotype indeed significantly limits the decrease of susceptible cells and the corresponding reduction of the environmental pool of OtV virions at the end of the ‘epidemic’ stage, which effectively annihilates the potential for a ‘recovery’ phase and oscillatory dynamics.

When the phenotypic switch to resistance is induced by viral contact (IPP), the outcomes of *O. tauri*-OtV interactions (with respect to the rate of switch towards resistance, *δ*_e_, *δ*_i_, and the individual cost of such resistance, *ζ*) are very similar to those described when antiviral resistance is associated to GM in Fig. 4a.1 and 4a.2 (Supporting Information Fig. S2). The interaction is predicted to produce ‘boom and bust’ dynamics with even sharper peaks of OtV virions in the ‘epidemic’ phase since the pool of resistant cells (slowing down the spread of the virus) has to build-up through infections and is therefore selected latter on during such a phase. Accordingly, no stable coexistence was ever predicted, whatever the conditions being considered, even in the presence of PCD. Meanwhile, when the resistant cells are allowed to switch back to a susceptible phenotype (*δ*^′^ > 0), the *O. tauri*-OtV interactions always lead to a stable coexistence (Supporting Information Fig. S2), in the same way as when phenotypic switches were assumed to occur randomly (see Fig. 4b.3,b.4). As described earlier, the switch back from resistant to susceptible cells indeed allows to prevent the strong decreases in the density of both *O. tauri* susceptible cells and virions.

## Discussion

Since its first report in a phytoplankton species (Waters and Chan 1982), many studies have confirmed the existence of antiviral resistance in prokaryotic and eukaryotic species (Thyrhaug et al. 2003; Tomaru et al. 2009; Kimura and Tomaru 2014; Zborowsky and Lindell 2019; Yau et al. 2018, 2020; Esmael et al. 2023; Bedi De Silva et al. 2024; Shaler et al. 2025). While -omics studies are starting to unravel the diversity of determinisms and mechanisms of such individual resistances (Yau et al. 2020; Shaler et al. 2025; Schleyer et al. 2023; Zborowsky et al. 2025; James et al. 2026), the effects of these alternative molecular determinants on the population dynamics of phytoplankton-virus interactions remain broadly unknown. We provide here the first comprehensive theoretical investigation of those impacts through the systematic analysis of 12 epidemiological models allowing to scale-up from a cell-centred molecular description of antiviral resistance to the phytoplankton-virus interactions dynamics. The tailoring of those models to key *Mamiellophyceae*-prasinoviruses interactions, between *O. tauri* and its lytic OtV, unravelled the quantitative impacts of antiviral resistance on the emergence of lytic viruses and their ability to regulate phytoplanktonic species through ‘boom and bust’ or stable dynamics. The main outcomes of this integrative modelling study clearly show that the molecular determinism of antiviral resistance can have significant impacts at all stages of the phytoplankton-virus population dynamics.

### Random phenotypic plasticity between S and R cells is costly in a virus-free environment

First, before virus emergence, the individual cost paid by resistant individuals is typically known to limit the growth of the host population (Winter et al. 2010). The general expressions of the phytoplankton intrinsic growth rate (table 2, 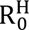), and their use to assess such an impact on *O. tauri*’s growth rate, further demonstrate that such a growth limitation strongly depends on the determinism of resistance. The effect on *O. tauri*’s growth rate is indeed obviously null when antiviral resistance is induced by the (absent) OtV, it reaches a maximum of 5% when resistance is associated to genomic mutations and rises to 20% when resistance is acquired through random phenotypic plasticity. Although those maximal effects were predicted for high individual costs of resistance, typically larger than the ≤1% thought to affect *Ostreoccocus* species (Thomas et al. 2011; Heath and Collins 2016; Heath et al. 2017), the ∼4.5-fold larger impact of randomly plastic strategy was consistently observed at such a lower cost. Those differences are in line with the basic understanding that, in the absence of viral selective pressures, genomic mutations provide less opportunities to sustain resistant individuals as they typically occur at lower rates (Yau et al. 2016, 2018; Krasovec et al. 2017). Meanwhile, the impacts associated with a random phenotypic plasticity of resistance were shown to be potentially higher on the 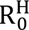 of *O. tauri* populations than the costs set at the individual level. Interestingly, this points out at a twofold effect of such a resistance determinism; beside the direct decrease of their rate of asexual reproduction (i.e. *b*’<*b* in table 1), resistant individuals incur a reduction of the period of time when cell division can occur that we show to be equal to (2*d*+*γ*+*γ*^′^)^-1^ (as compared to (2*d*)^-1^ when resistance is associated with genomic mutations). Such an unrecognized indirect cost could readily be a limiting factor as phenotypic switch to resistance has been shown to be associated with significant transcriptional changes (in *Ostreoccoccus* species (Yau et al. 2016, 2020), and other phytoplanktonic species (Shaler et al. 2025)) that are likely to impede the molecular initiation of cell division. Noteworthily, the above impacts on the phytoplankton intrinsic growth rate had little consequences on the actual abundance that can be reached by the corresponding populations. The effect of resistance on their biotic capacity was indeed shown to always remain <1.5%, whatever the determinism of antiviral resistance. Resistant cells paying the highest individual cost to resistance, and inducing those largest populational effects, are indeed unable to increase in frequencies larger than ∼10% in a virus-free environment where they are kept in check by intraspecific competition with susceptible ones.

### Random phenotypic plasticity provides a strong barrier to virus emergence

The early stage of the phytoplankton-virus interaction is also significantly affected by the determinism of antiviral resistance as it sets the conditions of virus emergence. Although the overall abundance of *O. tauri* is predicted to be always close to its maximal value in a virus free environment (see above), the percentage of resistant individuals in such populations was predicted to be ∼2-20 times higher when resistance is acquired through random phenotypic plasticity than when it is acquired through genomic mutations. As such differences are maximal for the low cost values typically associated with resistance in *Ostreoccocus* species (Thomas et al. 2011; Heath and Collins 2016; Heath et al. 2017), the determinism of antiviral resistance is indeed very likely to be influential in shaping the initial susceptibility of the phytoplankton population to the viral spread. Interestingly, the general expressions of the virus basic reproductive number (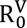 in table 3) unravelled that the percentage of reduction of OtV’ R_0_ merely matches the initial percentage of resistant cells. The most efficient barrier to virus emergence was therefore shown to be an antiviral resistance determined by random phenotypic plasticity, and its efficiency was even larger when such resistance was intracellular. In such a case, resistant cells represent ‘dead-end’ hosts, and they induce a strong ‘dilution effect’ (Keesing and Ostfeld 2021) decreasing OtV’ R_0_ by at least 50%, even at rate of phenotypic switches 10-100 times lower than those estimated in the closely related species *O. mediterraneus* (Yau et al. 2020). While the precise molecular mechanisms allowing *Ostreoccocus* species to resist remain to be fully characterized, there is evidence reported in *O. tauri* of an incomplete reduction of the rate of adsorption of virions on resistant cells (from 79% to 98%, (Yau et al. 2018)) consistent with an extracellular resistance mechanism.

Meanwhile, remnants of viral genome within the host genome (Blanc-Mathieu et al. 2017) and the externalization of virions by budding (Thomas et al. 2011) in *O. tauri* indicate that cells can also survive viral infection. Altogether, these suggest that antiviral resistance relies on an admixture of extracellular and intracellular strategies. According to our predictions, quantifying the relative importance of extracellular .*vs*. intracellular mechanisms of resistance is essential to assess its overall efficiency to limit the viral spread, especially at the low rates of random phenotypic plasticity occurring in *Ostreococcus* species (Yau et al. 2020).

### Bi-directional phenotypic plasticity stabilises phytoplankton-virus interactions

Third, once the virus has spread, the stability of the phytoplankton-virus dynamics strongly depends on antiviral resistance being acquired through either genomic mutations or phenotypic plasticity. In the former case, the interactions often lead to oscillatory dynamics, as expected from similar dynamics predicted by gene for gene modelling (Sasaki 2002) and the typically high burst-size associated with the lysis of infected phytoplankton cells, i.e. 25 virions in *O. tauri* (Derelle et al. 2008) and up to 330 in other *Mamiellophyceae* (*Micromonas pusilla* (Maat et al. 2016)). On the other hand, phenotypic switches between susceptible and resistant cells can readily allow for a stable equilibrium, as already suggested by a previous phenotypic model focused on the exponential growth of *O. mediterraneus* and its OmV virus in the early stage of experimental infections (Yau et al. 2020). The systematic analysis about the impact of both the rate of phenotypic acquisition and loss of resistance provided here clearly shows that, for both random and virus induced plasticity, the rate of phenotypic switches back to susceptible cells has a major impact on the stability of the interaction. This original prediction shed some light on the importance of the declining part of the ‘epidemic’ stage that follows the peak of viral infections in the stabilisation of *O. tauri*-OtV interaction dynamics. The phenotypic switches towards susceptibility then limit the crash of the susceptible part of the phytoplankton population and, thereby, of viruses, which leads to dampened oscillations. Such a stabilising effect is predicted to be strong enough to allow for a stable phytoplankton-virus equilibrium to be systematically reached.

The overall conclusion of this broad theoretical study on phytoplankton antiviral resistance is that random phenotypic plasticity provides the strongest barrier to viral emergence and promotes the stability of phytoplankton-virus interactions, although its maintenance in the absence of virus is obviously costly. Such a trade-off raises intriguing perspectives to think about the adaptive evolution of phytoplankton resistance in natural environments, whose levels of spatial heterogeneity remain broadly unknown at the incredibly small scale of their interactions, i.e. with densities up to 10^3^-10^4^ *Ostreoccocus* sp. (Demir-Hilton et al. 2011) and virions (Bellec et al. 2010) per ml. A heterogeneous distribution of virus could indeed result in random selective pressures that are well-known to select for random phenotypic plasticity providing efficient bet-hedging strategies (Yau et al. 2020). Meanwhile, the above trade-off may concomitantly lead to unravelled seasonal balanced selection regimes since viral pressures are expected to show marked infra-annual variations associated with variations of the mixed-layer depth ranging from tens to hundreds of metres (Moreles et al. 2025).

According to the well-recognized ‘Disturbance recovery’ hypothesis, such variations are indeed thought to induce essential variations in the density of zooplankton (and therefore of the ‘top-down’ control of phytoplankton species) shaping typical bloom dynamics (Behrenfeld and Boss 2014). As this study contributed to demonstrate, the molecular determinism of antiviral resistance truly affects the efficiency of the ‘top-down’ control exerted by lytic viruses, which in turn is likely to have an impact on phytoplankton diversity as virus-mediated coexistence is thought to be essential to explain phytoplankton diversity and solve the so-called ‘Paradox of the plankton’ (Flynn et al. 2022). This strongly points toward the need for integrative multi-omics approaches combined with eco-genomic modelling approach to improve our quantitative understanding of the dynamics of marine ecosystem and the services they provide.

## Author Contributions

Raphaël Rousseau: conceptualization, data curation, methodology, investigation, formal analysis, visualization, writing (original draft). Carlos Cáceres: methodology, writing (review & editing). Gwenaël Piganeau: funding acquisition, writing (review & editing). Sébastien Gourbière: conceptualization, methodology, funding acquisition, supervision, validation, visualization, writing (original draft).

## Acknowledgments

This study is set within the framework of the ‘Laboratoire d’Excellence (LABEX)’ TULIP (ANR-10-LABX-0041) and of the ‘École Universitaire de Recherche (EUR)’ TULIP-GS (ANR-18-EURE-0019). This work has benefited from a PhD fellowship to RR (Région Occitanie and TULIP-GS, ANR-18-EURE-0019) and from the project ‘Modélisation ‘Epidémio-Génomique’ des interactions algues-virus’ (MITI CNRS, PI. Gourbiere S.). This work further benefited from the modelling discussions hold in the context of the ‘PHYTOMICS’ (ANR, ANR-21-CE02-0026, PI. Gourbiere S.) and ‘ELVIRA’ (ANR, ANR-21-CE20-0041, PI. Piganeau G.) projects. The funders played no role in the study design, data collection and analysis, decision to publish, or preparation of the manuscript.

## Data Availability Statement

The data that support the findings of this study are available from the corresponding author upon request.

