## Supporting Information for "How do the molecular determinants of antiviral resistance shape the dynamic of phytoplankton-virus interaction?"

### Supporting Text

#### Heading 1. Detailed explanation of the estimates presented in tables 1 (main text) and S1.

**Table S1. Estimates of the SIV-SRIV modelling parameters describing *O. tauri*-OtV interactions.** The definition of all modelling parameters describing *O. tauri* and OtV life-history traits, their encounter rates, as well as the determinism and mechanisms of antiviral resistance are provided with the corresponding symbols, values, and units.

| Symbol | Definition | Value(s) | Unit | Reference |
|---|---|---|---|---|
| $b$ | asexual reproduction rate of susceptible hosts | 0.764 | $d^{-1}$ | Krasovec et al. 2016 (1) |
| $d$ | intrinsic mortality rate of host | 0.05 | $d^{-1}$ | Listmann et al. 2021 |
| $b'$ | asexual reproduction rate of resistant hosts | 0.688; 0.756; 0.763 | $d^{-1}$ | this study (2) |
| $c$ | intraspecific competition rate | $5\times10^{-7}$ | $d^{-1}$ | this study (3) |
| $\mu$ | probability of mutation (S → R) | $10^{-10}$ to $10^{-2}$ | $division^{-1}$ | Krasovec et al. 2017 (4) |
| $\gamma$ | rate of random phenotypic plasticity (S → R) | $10^{-4}$ to $10^{-2}$ | $d^{-1}$ | Yau et al. 2018 (4) |
| $\delta_e, \delta_i$ | rate of induced phenotypic plasticity (S → R) | $10^{-4}$ to $10^{-2}$ | $d^{-1}$ | Yau et al. 2018 (4) |
| $\gamma', \delta'$ | rates of phenotypic plasticity (R → S) | 0 to $10^{-2}$ | $d^{-1}$ | this study |
| $\beta$ | infectious encounter rate of susceptible hosts | $1.37\times10^{-8}$ to $1.37\times10^{-6}$ | $ml.d^{-1}$ | this study (5) |
| $\beta'$ | encounter rate of resistant hosts | 0 or $\beta$ | $ml.d^{-1}$ | this study |
| $\alpha$ | lysis rate | 2.5 | $d^{-1}$ | Derelle et al. 2008; Monier et al. 2017; Derelle et al. 2018 (6) |
| $b_v$ | burst size | 25 | $virions.cell^{-1}$ | Derelle et al. 2008 |
| $d_v$ | decay rate of virions | 0.01 | $d^{-1}$ | Listmann et al. 2021 |
| $\alpha'$ | rate of PCD | 0 to $2\alpha$ | $d^{-1}$ | this study (7) |

(1) The population growth rate of *O. tauri* ($r = b - d$) was estimated by fitting an exponential model to experimental kinetics in (Krasovec et al. 2016). The rate of asexual reproduction ($b$) was then calculated by summing the population growth rate ($r$) and the mortality rate ($d$) estimated respectively in (Krasovec et al. 2016) and (Listmann et al. 2021). (2) While the cost of antiviral resistance is thought to be low in *O. tauri* populations (Heath and Collins 2016; Heath et al. 2017), and yet susceptible cells grow faster than resistant ones (Thomas et al. 2011), experimental studies remain to be set-up to provide proper estimates of such a cost. We therefore considered costs ($\zeta$) of 0.1%, 1% and 10%, the upper limit being set from a population dynamic study on cyanobacteria, another key group of unicellular phytoplankton (Lennon et al. 2007).

The rate of asexual reproduction of resistant cells was then set to $b' = b(1 - \zeta)$. (3) The intensity of intraspecific competition ($c$) was derived from the expression of the phytoplankton biotic capacity obtained from our core SIV model in the absence of virus, i.e. $K_N = r/c$, and by using both our estimate of the population growth rate ($r$) and the maximal density observed in experimental cultures of *O. tauri* (Thomas et al. 2011; Heath and Collins 2016; Heath et al. 2017; Vacant et al. 2022; Barton et al. 2023). We then set $c$ to the average of the obtained values ($8.4\times10^{-9}$, $9.3\times10^{-7}$, $1.2\times10^{-6}$, $8.5\times10^{-8}$, $3.2\times10^{-7}$). (4) The investigated ranges of probability of mutation ($\mu$) and rates of phenotypic plasticity towards resistance ($\gamma$, $\delta_e$ and $\delta_i$) were set according to previous studies of the rate of spontaneous mutations (Krasovec et al. 2017) and phenotypic switches (Yau et al. 2018). (5) We estimated the encounter rate between *O. tauri* and its OtV using the well-recognised relationship proposed by Murray and Jackson (1992, (Murray and Jackson 1992)) that accounts for the size (diameter) of the host and virus in addition to parameters describing the abiotic conditions of the environment, i.e. temperature and salinity (see Supporting Information, Heading 5). Since this relationship typically provides maximum rates of encounter (Murray and Jackson 1992), we made predictions for rates ranging from such a maximum to a 1/100 of its value, the lower limit being set according to independent estimates derived from unpublished experimental data on *O. tauri*-OtV interaction dynamics. (6) The estimate of the lysis rate was taken as the average of its previous estimates. (7) The rate of PCD ($\alpha'$) was set to 0 (no PCD) and to values equal to once or twice the rate of lysis ($\alpha$) since such a collective defence mechanism only makes sense if the ‘altruistic suicide’ occurs before infection has led to the production of virions.

**Heading 2. Local stability analysis of the SRIV-GM model of phytoplankton-virus interaction with antiviral resistance associated to genomic mutation.**

**1. The SRIV-GM model**. As described in section 'Genomic mutation for antiviral resistance (GM)', the SRIV-GM model is defined from the following set of ordinary differential equations;

$$\frac{\mathrm{dS}}{\mathrm{d}t} = (b - c\mathrm{N})(1 - \mu)\mathrm{S} - dS - \beta\mathrm{SV} \qquad (\mathrm{S2}.a)$$

$$\frac{\mathrm{dR}}{\mathrm{d}t} = (b - c\mathrm{N})\mu S + (b' - c\mathrm{N})\mathrm{R} - d\mathrm{R} \qquad (\mathrm{S2}.b)$$

$$\frac{\mathrm{dI}}{\mathrm{d}t} = \beta\mathrm{SV} - (\alpha + \alpha' + d)\mathrm{I} \qquad (\mathrm{S2}.c)$$

$$\frac{\mathrm{dV}}{\mathrm{d}t} = b_v\alpha\mathrm{I} - d_v\mathrm{V} - \beta\mathrm{SV} - \beta'\mathrm{RV} \qquad (\mathrm{S2}.d)$$

where S, R, I and N stand for the density of susceptible, resistant, infected and total (N=S+R+I) phytoplanktonic cells, while V represents the density of virions in the marine environment.

**2. Stability analysis of the SRIV-GM model.** The SRIV-GM model has four fixed points; a trivial equilibrium where neither phytoplankton nor viruses persist, i.e. TE(0, 0, 0, 0), two Disease Free Equilibrium where either i) only resistant phytoplanktonic cells are present, i.e. $\mathrm{DFE}_1$ (0, $R^*$, 0, 0), or ii) susceptible and resistant phytoplanktonic cells persist, i.e. $\mathrm{DFE}_2$ ($S^*$, $R^*$, 0, 0), and a Disease Equilibrium where viruses persist and are found infecting the phytoplankton population, i.e. DE ($S^*$, $R^*$, $I^*$, $V^*$). The formal expressions of these four fixed points are provided below, while analysing their local stability properties.

The general expression of the Jacobian matrix that we used to assess the local stability of the four fixed points can be written as follows:

$$J(\mathrm{S}^*,\mathrm{R}^*,\mathrm{I}^*,\mathrm{V}^*) = \begin{pmatrix} \big(b - c(2\mathrm{S}^* + \mathrm{R}^* + \mathrm{I}^*)\big)(1-\mu) - d - \beta\mathrm{V}^* & -c(1-\mu)\mathrm{S}^* & -c(1-\mu)\mathrm{S}^* & -\beta\mathrm{S}^* \\ \big(b - c(2\mathrm{S}^* + \mathrm{R}^* + \mathrm{I}^*)\big)\mu - c\mathrm{R}^* & b' - c(\mathrm{S}^* + 2\mathrm{R}^* + \mathrm{I}^*) - d - c\mu\mathrm{S}^* & -c(\mu\mathrm{S}^* + \mathrm{R}^*) & 0 \\ \beta\mathrm{V}^* & 0 & -(\alpha + \alpha' + d) & \beta\mathrm{S}^* \\ -\beta\mathrm{V}^* & -\beta'\mathrm{V}^* & b_v\alpha & -d_v - \beta\mathrm{S}^* - \beta'\mathrm{R}^* \end{pmatrix}$$

where $b' = b(1-\zeta)$, and $\mathrm{S}^*$, $\mathrm{R}^*$, $\mathrm{I}^*$, $\mathrm{V}^*$ stand for the equilibrium density of (susceptible, resistant and infected) phytoplanktonic cells and virions (at any of the four fixed points evocated above).

*2.1. Stability analysis of the trivial equilibrium.* At the TE (0, 0, 0, 0), the above Jacobian matrix becomes:

$$J(0,0,0,0) = \begin{pmatrix} b(1-\mu) - d & 0 & 0 & 0 \\ b\mu & b' - d & 0 & 0 \\ 0 & 0 & -(\alpha + \alpha' + d) & 0 \\ 0 & 0 & b_v\alpha & -d_v \end{pmatrix}$$

Since $J(0,0,0,0)$ is block diagonal, its eigenvalues correspond to those of it two $2 \times 2$ diagonal submatrices. According to the Routh-Hurwitz criteria, the eigenvalues of those submatrices have negative real parts whenever their trace is negative and their determinant is positive (Otto and Day 2011).

It is straightforward to show that the lower diagonal submatrix satisfies those criteria, so that the stability of the trivial equilibrium ultimately depends on the eigenvalues of the upper diagonal submatrix. Its trace and determinant are equal to $b(1-\mu) + b' - 2d$ and $(b(1-\mu) - d)(b' - d)$, respectively. The determinant is positive since i) the intrinsic growth rates of susceptible and resistant individuals, i.e. $b - d$ and $b' - d$, are positive, and ii) the mutation probability ($\mu$) is typically much smaller than 1. Meanwhile, for the trace to be negative, the following condition has to be fulfilled;

$$\frac{b(1-\mu) + b'}{2d} < 1 \qquad \text{(S3)}$$

The trivial equilibrium is then stable whenever this condition is satisfied and, otherwise, the phytoplankton population is predicted to grow. Accordingly, the left-hand side of the above inequality corresponds to the intrinsic growth rate of a phytoplankton population ($\mathrm{R_0^H}$) appearing in table 2.

*2.2. Stability analysis of the resistant only DFE.* At the DFE$_1$ $(0, \mathrm{R}^*, 0,0)$, the expression of $\mathrm{R}^*$ simply reads;

$$\mathrm{R}^* = \frac{b' - d}{c} = \mathrm{K_N} \tag{S4}$$

The general expression of the Jacobian matrix then simplifies to:

$$J(0, \mathrm{R}^*, 0,0) = \begin{pmatrix} (b\zeta + d)(1-\mu) - d & 0 & 0 & 0 \\ (b\zeta + d)\mu - (b' - d) & -(b' - d) & -(b' - d) & 0 \\ 0 & 0 & -(\alpha + \alpha' + d) & 0 \\ 0 & 0 & b_v\alpha & -d_v - \beta'\mathrm{K_N} \end{pmatrix}$$

Since $J(0, \mathrm{R}^*, 0,0)$ is a block triangular matrix, their eigenvalues correspond to those of its two $2 \times 2$ diagonal submatrices.

As for the TE, the local stability of the DFE$_1$ primarily depends on the eigenvalues of the upper diagonal submatrix, as the Routh-Hurwitz criteria (trace < 0 and determinant > 0) are obviously satisfied by the lower diagonal submatrix. Since the upper submatrix is lower triangular, its eigenvalues correspond to its diagonal entries, which must be negative for the DFE$_1$ to be stable. The second eigenvalue $-(b' - d)$ is negative since the resistant's growth rate $(b' - d)$ is set to be positive. Meanwhile, for the first eigenvalue $(b\zeta + d)(1 - \mu) - d$ to be negative, the following condition must be satisfied;

$$\frac{b(1-\mu)}{d} < \frac{\mu}{\zeta} \tag{S5}$$

Although mathematically feasible, this condition is actually very unlikely to be met as it requires both a large mutation rate and a very low cost of resistance for the resistant cells to outcompete the susceptible ones.

*2.3. Stability analysis of the susceptible-resistant DFE.* At the DFE$_2$ $(\mathrm{S}^*, \mathrm{R}^*, 0{,}0)$, the expressions of $\mathrm{S}^*$ and $\mathrm{R}^*$ are:

$$\mathrm{S}^* = \mathrm{K_N} - \mathrm{R}^* \tag{S6.a}$$

$$\mathrm{R}^* = \frac{d\mu}{b\zeta(1-\mu)}\mathrm{K_N} \tag{S6.b}$$

where $\mathrm{K_N} = \frac{1}{c}\left(b - \frac{d}{(1-\mu)}\right)$.

The general expression of the Jacobian matrix then simplifies to:

$$J(\mathrm{S}^*, \mathrm{R}^*, 0{,}0) = \begin{pmatrix} -c(1-\mu)\mathrm{S}^* & -c(1-\mu)\mathrm{S}^* & -c(1-\mu)\mathrm{S}^* & -\beta\mathrm{S}^* \\ \big(b - c(2\mathrm{S}^* + \mathrm{R}^*)\big)\mu - c\mathrm{R}^* & b' - c(\mathrm{S}^* + 2\mathrm{R}^*) - d - c\mu\mathrm{S}^* & -c(\mu\mathrm{S}^* + \mathrm{R}^*) & 0 \\ 0 & 0 & -(\alpha + \alpha' + d) & \beta\mathrm{S}^* \\ 0 & 0 & b_v\alpha & -d_v - \beta\mathrm{S}^* - \beta'\mathrm{R}^* \end{pmatrix}$$

Since $J(\mathrm{S}^*, \mathrm{R}^*, 0{,}0)$ is a block triangular matrix, its eigenvalues correspond to those of their two $2 \times 2$ diagonal submatrices.

The trace and determinant of the upper diagonal submatrix are equal to $d\left(\frac{2}{1-\mu} - 1\right) - b(1+\zeta)$ and $cb\zeta(1-\mu)\mathrm{S}^*$, respectively. Given that i) $b > d$ so that the phytoplankton population can grow and ii) that $\mu$ is typically much smaller than 1, it is straightforward to show that the Routh-Hurwitz criteria (trace < 0 and determinant > 0) are satisfied.

Accordingly, the stability of the DFE ($\mathrm{S}^*$, $\mathrm{R}^*$, 0 ,0) ultimately depends on the eigenvalues of the lower diagonal submatrix of the above Jacobian. Although the trace of this submatrix is obviously negative, its determinant must be positive for the DFE to be stable, which led to the following condition;

$$\beta S^* \left(\frac{\alpha}{(\alpha + \alpha' + d)} b_v - 1\right) \frac{1}{d_v + \beta' R^*} < 1 \qquad (S7)$$

When such a condition is not verified, one expects the virus to be able to spread by infecting the susceptible individuals. Accordingly, its left-hand side represents the initial growth rate of the viral population, i.e. the virus basic reproduction number $\left(R_0^V\right)$ appearing in table 3.

*2.4. Stability analysis of the disease equilibrium.* At the DE ($S^*$, $R^*$, $I^*$, $V^*$), the formal expression of the densities of individuals can still be identified, although this requires the following convoluted calculations.

First, setting dI/dt = 0 allows for the identification of an expression for $I^*$. Second, the integration of this expression in the equilibrium condition dV/dt = 0 provides an expression for $S^*$. Third, an expression for $V^*$ is derived by combining the two previous expressions (for $I^*$ and $S^*$) with the equilibrium condition dS/dt = 0. Fourth, an expression for $R^*$ is obtained by incorporating all expressions for $S^*$, $I^*$ and $V^*$ into the last equilibrium condition dR/dt = 0. Since this last expression for $R^*$ only depends on the model parameters, it can, in turn, be used to make all other expressions (of $V^*$, $S^*$ and ultimately, $I^*$) explicit functions of those parameters.

By applying this analytical procedure to equations S2.*a*, S2.*c* and S2.*d*, we first expressed $S^*$, $I^*$ and $V^*$ as functions of $R^*$:

$$I^*(R^*) = A_1 S^*(R^*) V^*(R^*) \qquad (S8.a)$$

$$S^*(R^*) = A_2 + B_2 R^* \qquad (S8.b)$$

$$V^*(R^*) = \frac{A_3 + B_3 R^*}{\beta + c(1-\mu) A_1 (A_2 + B_2 R^*)} \qquad (S8.c)$$

where $A_1 = \frac{\beta}{(\alpha+\alpha'+d)}$, $A_2 = \frac{d_v}{(b_v\alpha A_1-\beta)}$, $B_2 = \frac{\beta'}{(b_v\alpha A_1-\beta)}$, $A_3 = (b - cA_2)(1-\mu) - d$ and $B_3 = -c(B_2+1)(1-\mu)$.

The three functions above were then substituted for $S^*$, $I^*$ and $V^*$ in equation S2.*b*, to derive the following cubic equation that allows for the identification of $R^*$ for the GM model:

$$f(R^*) = A_4 + B_4R^* + C_4R^{*2} + D_4R^{*3} = 0 \tag{S9}$$

where

$$A_4 = \beta(b - cA_2)\mu A_2 + c\big((b - cA_2)(1-\mu) - A_3\big)\mu A_1A_2^2$$

$$B_4 = b\mu(2c(1-\mu)A_1A_2 + \beta)B_2 + \beta(b' - d - cA_2) + c(b' - d - cA_2)(1-\mu)A_1A_2$$
$$-c\beta\mu A_2(2B_2+1) - c^2\mu(1-\mu)A_1A_2^2(3B_2+1) - c(2\mu B_2+1)A_1A_2A_3 - c\mu A_1A_2^2B_3$$

$$C_4 = c\big(bB_2 - cA_2(3B_2+2)\big)\mu(1-\mu)A_1B_2 + c\big((b'-d)B_2 - cA_2(2B_2+1)\big)(1-\mu)A_1$$
$$-c\beta\mu B_2(B_2+1) - c\beta(B_2+1) - cA_1A_2(2\mu B_2+1)B_3 - cA_1A_3B_2(\mu B_2+1)$$

$$D_4 = -c^2\mu(1-\mu)A_1B_2^2(B_2+1) - c^2(1-\mu)A_1B_2(B_2+1) - cA_1B_2(\mu B_2+1)B_3$$

While such an equation can be solved numerically, we looked for an approximate analytical expression of $R^*$ by perturbation analysis using Taylor series [75] in order to ease the subsequent identification of the conditions for the DE to be stable. This approximate expression of $R^*$ ($\widetilde{R^*}$; S10.*a*) was found to be the sum of a constant ($r_0$; S10.*b*) and a linear ($r_1$; S10.*c*) approximation whose (very impressive) accuracy is shown in Supporting Information, Heading 6.

$$\widetilde{R^*} = r_0 + r_1\mu \tag{S10.$a$}$$

$$r_0 = \frac{(b'-d)\beta\big(\alpha(b_v-1) - (\alpha'+d)\big) - c(b\zeta + \alpha + \alpha' + d)d_v}{c\Big(\beta'(b\zeta + \alpha + \alpha' + d) + \beta\big(\alpha(b_v-1) - (\alpha'+d)\big)\Big)} \tag{S10.$b$}$$

$$r_1 = \frac{r_{1A} + r_{1B}}{r_{1C}} \qquad (S10.c)$$

where

$$r_{1A} = \left(\frac{1}{r_0}\right)\left(b - c\frac{d_v(\alpha + \alpha')}{\beta\big(\alpha(b_v - 1) - (\alpha' + d)\big)}\right) + \beta'\frac{b}{d_v} + c\left(1 + \frac{\beta'}{d_v}r_0\right)\frac{b\zeta - (\alpha + \alpha')}{(\alpha + \alpha' + d)}$$

$$r_{1B} = -c\beta'\left(2 + \frac{\beta'}{d_v}r_0\right)\frac{(\alpha + \alpha')}{\beta\left(\alpha(b_v - 1) - (\alpha' + d)\right)}$$

$$r_{1C} = \frac{c\left(\beta'(b\zeta + \alpha + \alpha' + d) + \beta\big(\alpha(b_v - 1) - (\alpha' + d)\big)\right)}{d_v(\alpha + \alpha' + d)}$$

Using equation S10.*a*, we subsequently identified $S^*$, $I^*$ and $V^*$ according to the relationships S8.*a* to S8.*c*.

This allowed to perform a local stability of the DE by examining the Routh–Hurwitz criteria for a $4 \times 4$ matrix. To ensure that all eigenvalues of the matrix have a negative real part, the followings inequality are satisfied (Otto and Day 2011):

$$a_1 > 0 \qquad (S11.a)$$

$$a_3 > 0 \qquad (S11.b)$$

$$a_4 > 0 \qquad (S11.c)$$

$$a_1 a_2 a_3 > a_3^2 + a_1^2 a_4 \qquad (S11.d)$$

where $a_1$ to $a_4$ denote the coefficients of the characteristic polynomial of the Jacobian:

$$\lambda^4 + a_1\lambda^3 + a_2\lambda^2 + a_3\lambda + a_4 \qquad (S12)$$

For the Jacobian matrix $J(S^*, R^*, I^*, V^*)$ associated with the DE of our SRIV-GM model, the expressions of those coefficients are the following:

$$a_1 = (\alpha + \alpha' + 2d) + \beta b_v \frac{\alpha}{(\alpha + \alpha' + d)} \mathrm{S}^* + c(2\mathrm{S}^* + 2\mathrm{R}^* + \mathrm{I}^*) - b'$$

$$a_2 = \big(c(2\mathrm{S}^* + 2\mathrm{R}^* + \mathrm{I}^*) - (b' - d)\big)\left(\beta b_v \frac{\alpha}{(\alpha + \alpha' + d)} \mathrm{S}^* + (\alpha + \alpha' + d)\right) + c\beta(1-\mu)\mathrm{S}^*\mathrm{V}^*$$

$$+c\big(b - c(\mathrm{S}^* + \mathrm{R}^* + \mathrm{I}^*)\big)\mu(1-\mu)\mathrm{S}^* - \beta^2\mathrm{S}^*\mathrm{V}^* - c\big(b' - d - c(\mathrm{S}^* + \mathrm{R}^* + \mathrm{I}^*)\big)(1-\mu)\mathrm{S}^*$$

$$a_3 = \big(b - c(\mathrm{S}^* + \mathrm{R}^* + \mathrm{I}^*)\big)\mu\left(c\beta b_v \frac{\alpha}{(\alpha + \alpha' + d)}(1-\mu)\mathrm{S}^{*2} + c(\alpha + \alpha' + d)(1-\mu)\mathrm{S}^*\right)$$

$$+\beta^2 b_v \alpha \mathrm{S}^*\mathrm{V}^* + c\beta^2 b_v \frac{\alpha}{(\alpha + \alpha' + d)}(1-\mu)\mathrm{S}^{*2}\mathrm{V}^* - \beta^2(\alpha + \alpha' + d)\mathrm{S}^*\mathrm{V}^*$$

$$-\big(b' - d - c(\mathrm{S}^* + \mathrm{R}^* + \mathrm{I}^*)\big)\left(c\beta b_v \frac{\alpha}{(\alpha + \alpha' + d)}(1-\mu)\mathrm{S}^{*2} + c(\alpha + \alpha' + d)(1-\mu)\mathrm{S}^*\right.$$

$$\left.+c\beta(1-\mu)\mathrm{S}^*\mathrm{V}^* - \beta^2\mathrm{S}^*\mathrm{V}^*\right) - c\beta^2(\mathrm{S}^* + \mathrm{R}^*)\mathrm{S}^*\mathrm{V}^* - \beta\beta'\big(b - c(\mathrm{S}^* + \mathrm{R}^* + \mathrm{I}^*)\big)\mu\mathrm{S}^*\mathrm{V}^*$$

$$a_4 = c(\mu\mathrm{S}^* + \mathrm{R}^*)(\beta^2\beta'\mathrm{S}^*\mathrm{V}^{*2} + \beta^2 b_v \alpha \mathrm{S}^*\mathrm{V}^* + \beta\beta'(\alpha + \alpha' + d)\mathrm{S}^*\mathrm{V}^* - \beta^2(\alpha + \alpha' + d)\mathrm{S}^*\mathrm{V}^*)$$

$$+\big(b' - d - c(\mathrm{S}^* + \mathrm{R}^* + \mathrm{I}^*)\big)\left(\beta^2(\alpha + \alpha' + d)\mathrm{S}^*\mathrm{V}^* + c\beta^2(1-\mu)\mathrm{S}^{*2}\mathrm{V}^* - \beta^2 b_v \alpha \mathrm{S}^*\mathrm{V}^*\right.$$

$$\left.-c\beta^2 b_v \frac{\alpha}{(\alpha + \alpha' + d)}(1-\mu)\mathrm{S}^{*2}\mathrm{V}^*\right) - (\beta\beta'(\alpha + \alpha' + d)\mathrm{S}^*\mathrm{V}^* + c\beta\beta'(1-\mu)\mathrm{S}^{*2}\mathrm{V}^*)\mu \times$$

$$\big(b - c(\mathrm{S}^* + \mathrm{R}^* + \mathrm{I}^*)\big)$$

The above expressions, together with the expressions of $\mathrm{S}^*$, $\mathrm{R}^*$, $\mathrm{I}^*$ and $\mathrm{V}^*$ (S8*a*-*c* and S10*a*-*c*), were then used to assess numerically if the Routh–Hurwitz criteria were satisfied and therefore the DE locally stable.

**Heading 3. Local stability analysis of the SRIV-RPP model of phytoplankton-virus interaction with antiviral resistance associated to random phenotypic plasticity.**

**1. The SRIV-RPP model**. As described in section 'Phenotypic plasticity between susceptibility and antiviral resistance (PP)', the SRIV-RPP model is defined by the following set of ordinary differential equations;

$$\frac{\mathrm{dS}}{\mathrm{d}t} = (b - c\mathrm{N})\mathrm{S} - d\mathrm{S} + \gamma'\mathrm{R} - \gamma\mathrm{S} - \beta\mathrm{SV} \qquad (\mathrm{S}13.a)$$

$$\frac{\mathrm{dR}}{\mathrm{d}t} = (b' - c\mathrm{N})\mathrm{R} - d\mathrm{R} + \gamma\mathrm{S} - \gamma'\mathrm{R} \qquad (\mathrm{S}13.b)$$

$$\frac{\mathrm{dI}}{\mathrm{d}t} = \beta\mathrm{SV} - (\alpha + \alpha' + d)\mathrm{I} \qquad (\mathrm{S}13.c)$$

$$\frac{\mathrm{dV}}{\mathrm{d}t} = b_v\alpha\mathrm{I} - d_v\mathrm{V} - \beta\mathrm{SV} - \beta'\mathrm{RV} \qquad (\mathrm{S}13.d)$$

where S, R, I and N stand for the density of susceptible, resistant, infected and total (S+R+I) phytoplanktonic cells, while V represents the density of virions in the marine environment.

**2. Stability analysis of the SRIV-RPP model.** The above RPP model has three fixed points; a trivial equilibrium TE(0, 0, 0, 0) where neither phytoplankton nor viruses persist, a Disease Free Equilibrium where susceptible and resistant cells can persist since phenotypic switches between S and R individuals occur even in the absence of virus, i.e. DFE ($S^*$, $R^*$, 0, 0), and a Disease Equilibrium DE ($S^*$, $R^*$, $I^*$, $V^*$) where viruses persist and infect the phytoplankton population. The formal expressions of these three fixed points are provided below, while analysing their local stability properties.

The general expression of the Jacobian matrix, that we subsequently used to assess the local stability of these three fixed points, can be shown to be:

$$J(\mathrm{S}^*,\mathrm{R}^*,\mathrm{I}^*,\mathrm{V}^*) = \begin{pmatrix} r - c(2\mathrm{S}^* + \mathrm{R}^* + \mathrm{I}^*) - \gamma - \beta\mathrm{V}^* & \gamma' - c\mathrm{S}^* & -c\mathrm{S}^* & -\beta\mathrm{S}^* \\ \gamma - c\mathrm{R}^* & b' - c(\mathrm{S}^* + 2\mathrm{R}^* + \mathrm{I}^*) - d - \gamma' & -c\mathrm{R}^* & 0 \\ \beta\mathrm{V}^* & 0 & -(\alpha + \alpha' + d) & \beta\mathrm{S}^* \\ -\beta\mathrm{V}^* & -\beta'\mathrm{V}^* & b_v\alpha & -d_v - \beta\mathrm{S}^* - \beta'\mathrm{R}^* \end{pmatrix}$$

where $b' = b(1-\zeta)$, and $\mathrm{S}^*$, $\mathrm{R}^*$, $\mathrm{I}^*$, $\mathrm{V}^*$ stand for the number of (susceptible, resistant and infected) phytoplanktonic cells and virions (at any of the three fixed points evocated above).

*2.1. Stability analysis of the trivial equilibrium.* At the TE $(0,0,0,0)$, the above Jacobian become:

$$J(0,0,0,0) = \begin{pmatrix} b - d - \gamma & \gamma' & 0 & 0 \\ \gamma & b' - d - \gamma' & 0 & 0 \\ 0 & 0 & -(\alpha + \alpha' + d) & 0 \\ 0 & 0 & b_v\alpha & -d_v \end{pmatrix}$$

Since $J(0,0,0,0)$ is a block diagonal matrix, its eigenvalues correspond to those of its two diagonal submatrices. According to the Routh-Hurwitz criteria, the equilibrium is stable whenever the eigenvalues of those submatrices have negative real parts (Otto and Day 2011).

It is straightforward to show that the lower diagonal submatrix of $J(0,0,0,0)$ satisfies those criteria, so that the stability of the trivial equilibrium only depends on the eigenvalues of the upper diagonal submatrix. Its trace and determinant are equal to $b + b' - 2d - \gamma - \gamma'$ and $(b-d)(b'-d) - (b-d)\gamma' - (b'-d)\gamma$, respectively. For the determinant to be positive, the following condition has to be fulfilled;

$$\frac{\gamma}{b-d} + \frac{\gamma'}{b'-d} < 1 \tag{S14}$$

This is always the case since i) the intrinsic growth rates of susceptible and resistant individuals, i.e. $b - d$ and $b' - d$, are of the same order as 1, whereas the rates of phenotypic switches $\gamma$

(from S to R) and $\gamma'$ (from R to S) are typically much lower than 1. Meanwhile, for the trace to be negative, the following condition has to be fulfilled;

$$\frac{b+b'}{2d+\gamma+\gamma'}<1 \tag{S15}$$

The trivial equilibrium is then stable when this condition is satisfied and, otherwise, the phytoplankton population is predicted to grow. Accordingly, the left-hand side of the above inequality corresponds to the intrinsic growth rate of a phytoplankton population ($\mathrm{R}_0^{\mathrm{H}}$) appearing in table 2.

*2.2. Stability analysis of the disease-free equilibrium*. At the DFE $(\mathrm{S}^*, \mathrm{R}^*, 0, 0)$, the expression of $\mathrm{S}^*$ and $\mathrm{R}^*$ depends on the biotic capacity of phytoplankton ($\mathrm{K_N}$). To identify $\mathrm{K_N}$, we used the conditions corresponding to S13.*a* and S13.*b* at the DFE, which led to the following equalities where $r = b - d$ and $r' = b' - d$;

$$\frac{\mathrm{R}^*}{\mathrm{S}^*}=\frac{(c\mathrm{K_N}+\gamma-r)}{\gamma'} \tag{S16.$a$}$$

$$\frac{\mathrm{R}^*}{\mathrm{S}^*}=\frac{\gamma}{(c\mathrm{K_N}+\gamma'-r')} \tag{S16.$b$}$$

Combining these two equalities provided a quadratic equation:

$$f(\mathrm{K_N})=c^2\mathrm{K_N^2}+c(\gamma+\gamma'-r-r')\mathrm{K_N}+rr'-r'\gamma-r\gamma'=0 \tag{S17}$$

whose two roots are given by;

$$\mathrm{K_N}=\frac{r+r'-\gamma-\gamma'-\sqrt{(r-r')^2+(\gamma+\gamma')^2+2(r-r')(\gamma'-\gamma)}}{2c} \tag{S18.$a$}$$

$$\mathrm{K_N}=\frac{r+r'-\gamma-\gamma'+\sqrt{(r-r')^2+(\gamma+\gamma')^2+2(r-r')(\gamma'-\gamma)}}{2c} \tag{S18.$b$}$$

Numerical investigations showed that, in standard conditions (i.e. $\gamma$ and $\gamma' > 0$), the unique positive solution corresponds to the largest of the two roots (S18.$b$) that allowed to complete the expressions of $S^*$ and $R^*$ as follows:

$$S^* = R^* - K_N \tag{S19.a}$$

$$R^* = \frac{\gamma}{(cK_N - r' + \gamma + \gamma')} K_N \tag{S19.b}$$

At the DFE, the general expression of the Jacobian matrix simplifies to:

$$J(S^*, R^*, 0,0) = \begin{pmatrix} -\gamma'\frac{R^*}{S^*} - cS^* & \gamma' - cS^* & -cS^* & -\beta S^* \\ \gamma - cR^* & -\gamma\frac{S^*}{R^*} - cR^* & -cR^* & 0 \\ 0 & 0 & -(\alpha + \alpha' + d) & \beta S^* \\ 0 & 0 & b_v\alpha & -d_v - \beta S^* - \beta' R^* \end{pmatrix}$$

Since $J(S^*, R^*, 0,0)$ is a block triangular matrix, its eigenvalues correspond to those of the two diagonal submatrices.

The trace and determinant of the upper diagonal are equal to $-\gamma'\frac{R^*}{S^*} - \gamma\frac{S^*}{R^*} - cK_N$ and $c\left(1 + \frac{R^*}{S^*}\right)\gamma' R^* + c\left(1 + \frac{S^*}{R^*}\right)\gamma S^*$, respectively. Given that $S^*$and $R^*$ are both positive, the Routh-Hurwitz criteria are satisfied for this submatrix. Accordingly, the stability of the DFE ($S^*$, $R^*$,0,0) relies on the eigenvalues of the lower diagonal submatrix of J($S^*$, $R^*$,0, 0). Although the trace of this submatrix is obviously negative, its determinants must be positive for the DFE to be stable, which leads to the following condition;

$$\beta S^* \left(\frac{\alpha}{(\alpha + \alpha' + d)} b_v - 1\right) \frac{1}{d_v + \beta' R^*} < 1 \tag{S20}$$

When such a condition is not verified, one expects the virus to be able to spread by infecting the susceptible individuals. Accordingly, its left-hand side represents the initial growth rate of the viral population, i.e. the virus basic reproduction number $\left(R_0^V\right)$ appearing in table 3.

*2.3. Stability analysis of the disease equilibrium.* At the DE ($S^*$, $R^*$, $I^*$, $V^*$), the formal expression of the densities of individuals can still be identified, although this requires the same convoluted calculations as described in the analysis of the SRIV-GM model (Supporting Information, Heading 2). By applying the same process to equations S13.*a*, S13.*c* and S13.*d*, we expressed $S^*$, $I^*$ and $V^*$ as functions of $R^*$;

$$I^*(R^*) = A_1 S^*(R^*) V^*(R^*) \qquad (S21.a)$$

$$S^*(R^*) = A_2 + B_2 R^* \qquad (S21.b)$$

$$V^*(R^*) = \frac{A_3 + B_3 R^*}{\beta + c A_1 (A_2 + B_2 R^*)} \qquad (S21.c)$$

where $A_1 = \frac{\beta}{(\alpha+\alpha'+d)}$, $A_2 = \frac{d_v}{(b_v \alpha A_1 - \beta)}$, $B_2 = \frac{\beta'}{(b_v \alpha A_1 - \beta)}$, $A_3 = b - d - cA_2 - \gamma$ and $B_3 = \frac{\gamma'}{(A_2 + B_2 R^*)} - c(B_2 + 1)$.

The three functions above were then substituted for $S^*$, $I^*$ and $V^*$ in equation S13.*b*, to derive a quadratic equation allowing for the identification of $R^*$:

$$f_1(R^*) = A_4 + B_4 R^* + C_4 R^{*2} = 0 \qquad (S22)$$

Where $A_4 = \beta\gamma A_2 + c\gamma A_1 A_2^2$, $B_4 = \beta(b' - d - cA_2 - \gamma') + \beta\gamma B_2 + c\gamma A_1 A_2 (2B_2 + 1) - c(b\zeta + \gamma') A_1 A_2$ and $C_4 = c(\gamma - \gamma' - b\zeta) A_1 B_2 + c\gamma A_1 B_2^2 - c\beta(B_2 + 1) - c\gamma' A_1$.

The equation (S22) has only one positive solution providing the following expression of $R^*$:

$$R^* = \frac{-B_4 - \sqrt{B_4^2 - 4A_4C_4}}{2C_4} \qquad (S23)$$

Equation S23 was subsequently used to identify $S^*$, $I^*$ and $V^*$ according to the relationships S21.*a* to S21.*c*.

The conditions for the DE to be stable are then given by the Routh–Hurwitz criteria for a 4 × 4 matrix (Otto and Day 2011):

$$a_1 > 0 \quad (S24.a)$$

$$a_3 > 0 \quad (S24.b)$$

$$a_4 > 0 \quad (S24.c)$$

$$a_1 a_2 a_3 > a_3^2 + a_1^2 a_4 \quad (S24.d)$$

where $a_1$ to $a_4$ denote the coefficients of the characteristic polynomial of the Jacobian:

$$\lambda^4 + a_1\lambda^3 + a_2\lambda^2 + a_3\lambda + a_4 \quad (S25)$$

For the Jacobian matrix associated with the RPP model, these coefficients are given by:

$$a_1 = \beta b_v \frac{\alpha}{(\alpha + \alpha' + d)} \mathrm{S}^* + (\alpha + \alpha' + d) + c\mathrm{S}^* + c\mathrm{R}^* + \gamma \frac{\mathrm{S}^*}{\mathrm{R}^*} + \gamma' \frac{\mathrm{R}^*}{\mathrm{S}^*}$$

$$a_2 = \gamma'(\alpha + \alpha' + d)\frac{\mathrm{R}^*}{\mathrm{S}^*} + c(\alpha + \alpha' + d)\mathrm{S}^* + c\beta\mathrm{S}^*\mathrm{V}^* + \beta b_v \frac{\alpha}{(\alpha + \alpha' + d)} p\mathrm{R}^*$$

$$+c\beta b_v \frac{\alpha}{(\alpha + \alpha' + d)} \mathrm{S}^{*2} + c\mathrm{R}^* \left(\gamma' \frac{\mathrm{R}^*}{\mathrm{S}^*} + \beta b_v \frac{\alpha}{(\alpha + \alpha' + d)} \mathrm{S}^* + (\alpha + \alpha' + d) + \gamma'\right)$$

$$+\gamma \frac{\mathrm{S}^*}{\mathrm{R}^*}\left(c\mathrm{S}^* + \beta b_v \frac{\alpha}{(\alpha + \alpha' + d)} \mathrm{S}^* + (\alpha + \alpha' + d)\right) + c\gamma\mathrm{S}^* - \beta^2\mathrm{S}^*\mathrm{V}^*$$

$$a_3 = \beta^2 b_v \alpha \mathrm{S}^*\mathrm{V}^* + c(\alpha + \alpha' + d)\gamma\mathrm{S}^* + c\beta^2 b_v \frac{\alpha}{(\alpha + \alpha' + d)} \mathrm{S}^{*2}\mathrm{V}^* + c\beta b_v \frac{\alpha}{(\alpha + \alpha' + d)} \gamma\mathrm{S}^{*2}$$

$$+\gamma \frac{\mathrm{S}^*}{\mathrm{R}^*}\left(c\beta\mathrm{S}^*\mathrm{V}^* + c(\alpha + \alpha' + d)\mathrm{S}^* + c\beta b_v \frac{\alpha}{(\alpha + \alpha' + d)} \mathrm{S}^{*2} - \beta^2\mathrm{S}^*\mathrm{V}^*\right)$$

$$+c\mathrm{R}^*\left((\alpha + \alpha' + d)\gamma' + \beta b_v \frac{\alpha}{(\alpha + \alpha' + d)} \gamma'\mathrm{S}^* + \beta p\mathrm{V}^* + \gamma'(\alpha + \alpha' + d)\frac{\mathrm{R}^*}{\mathrm{S}^*}\right.$$

$$+\beta b_v \frac{\alpha}{(\alpha+\alpha'+d)}\gamma' \mathrm{R}^* - \beta^2 \mathrm{S}^*\mathrm{V}^*\Big) - c\beta^2 \mathrm{S}^{*2}\mathrm{V}^* - \beta^2(\alpha+\alpha'+d)\mathrm{S}^*\mathrm{V}^* - \beta\beta'\gamma \mathrm{S}^*\mathrm{V}^*$$

$$a_4 = \gamma \frac{\mathrm{S}^*}{\mathrm{R}^*}\left(\beta^2 b_v \alpha \mathrm{S}^*\mathrm{V}^* + c\beta^2 b_v \frac{\alpha}{(\alpha+\alpha'+d)}\mathrm{S}^{*2}\mathrm{V}^* - c\beta^2\mathrm{S}^{*2}\mathrm{V}^* - \beta^2(\alpha+\alpha'+d)\mathrm{S}^*\mathrm{V}^*\right)$$

$$+c\mathrm{R}^*\Big(\beta^2\beta'\mathrm{S}^*\mathrm{V}^{*2} + \beta^2 b_v \frac{\alpha}{(\alpha+\alpha'+d)}\gamma'\mathrm{S}^*\mathrm{V}^* + \beta^2 b_v \alpha \mathrm{S}^*\mathrm{V}^* + \beta\beta'(\alpha+\alpha'+d)\mathrm{S}^*\mathrm{V}^*$$

$$-\beta^2(\alpha+\alpha'+d)\mathrm{S}^*\mathrm{V}^* - \beta^2\gamma'\mathrm{S}^*\mathrm{V}^* - \beta\beta'\gamma'\mathrm{R}^*\mathrm{V}^*\Big) - \beta\beta'(\alpha+\alpha'+d)\gamma\mathrm{S}^*\mathrm{V}^* - c\beta'\beta\gamma\mathrm{S}^{*2}\mathrm{V}^*$$

Those expressions, together with the expressions of $\mathrm{S}^*$, $\mathrm{R}^*$, $\mathrm{I}^*$ and $\mathrm{V}^*$ (S21*a*-*c* and S23), were used to numerically assess the local stability of the DE.

**Heading 4. Local stability analysis of the SRIV-IPP model of phytoplankton-virus interaction with antiviral resistance associated to virus-induced phenotypic plasticity.**

**1. The SRIV-IPP models**. As described in section 'Phenotypic plasticity between susceptibility and antiviral resistance (PP)', the SRIV-IPP is defined by the following set of ordinary differential equations when considering extracellular resistance;

$$\frac{\mathrm{dS}}{\mathrm{d}t} = (b - c\mathrm{N})\mathrm{S} - d\mathrm{S} + \delta'\mathrm{R} - \beta\mathrm{SV} \qquad (S26.a)$$

$$\frac{\mathrm{dR}}{\mathrm{d}t} = (b' - c\mathrm{N})\mathrm{R} - d\mathrm{R} + \delta_{\mathrm{e}}\beta\mathrm{SV} - \delta'\mathrm{R} \qquad (S26.b)$$

$$\frac{\mathrm{dI}}{\mathrm{d}t} = (1 - \delta_{\mathrm{e}})\beta\mathrm{SV} - (\alpha + \alpha' + d)\mathrm{I} \qquad (S26.c)$$

$$\frac{\mathrm{dV}}{\mathrm{d}t} = b_v\alpha\mathrm{I} - d_v\mathrm{V} - (1 - \delta_{\mathrm{e}})\beta\mathrm{SV} \qquad (S26.d)$$

and to a second set of equations when resistance is assumed to be intracellular;

$$\frac{\mathrm{dS}}{\mathrm{d}t} = (b - c\mathrm{N})\mathrm{S} - d\mathrm{S} + \delta'\mathrm{R} - \beta\mathrm{SV} \qquad (S27.a)$$

$$\frac{\mathrm{dR}}{\mathrm{d}t} = (b' - c\mathrm{N})\mathrm{R} - d\mathrm{R} + \delta_{\mathrm{i}}\mathrm{I} - \delta'\mathrm{R} \qquad (S27.b)$$

$$\frac{\mathrm{dI}}{\mathrm{d}t} = \beta\mathrm{SV} - (\alpha + \alpha' + d + \delta_{\mathrm{i}})\mathrm{I} \qquad (S27.c)$$

$$\frac{\mathrm{dV}}{\mathrm{d}t} = b_v\alpha\mathrm{I} - d_v\mathrm{V} - \beta\mathrm{SV} - \beta'\mathrm{RV} \qquad (S27.d)$$

where S, R, I and N stand for the density of susceptible, resistant, infected and total (S+R+I) phytoplanktonic cells, while V represents the density of virions in the marine environment.

**2. Stability analysis of the IPP models.** The two above models have three fixed points; a trivial equilibrium TE (0, 0, 0, 0) where neither phytoplankton nor viruses persist, a Disease Free Equilibrium DFE ($S^*$, 0, 0, 0) where only susceptible cells are present, and a Disease Equilibrium DE ($S^*$, $R^*$, $I^*$, $V^*$) where viruses persist and infect the phytoplankton population. The formal expressions of these three fixed points are provided below, while analyzing their local stability properties.

The general expression of the Jacobian matrix, that we subsequently used to assess the local stability of these three fixed points assuming an extracellular resistance, can be shown to be:

$$J_1(S^*,R^*,I^*,V^*) = \begin{pmatrix} b - c(2S^* + R^* + I^*) - d - \beta V^* & \delta' - cS^* & -cS^* & -\beta S^* \\ \delta_e \beta V^* - cR^* & b' - c(S^* + 2R^* + I^*) - d - \delta' & -cR^* & \delta_e \beta S^* \\ (1-\delta_e)\beta V^* & 0 & -(\alpha + \alpha' + d) & (1-\delta_e)\beta S^* \\ -(1-\delta_e)\beta V^* & 0 & b_v \alpha & -d_v - (1-\delta_e)\beta S^* \end{pmatrix}$$

while the Jacobian matrix assuming an intracellular resistance is:

$$J_2(S^*,R^*,I^*,V^*) = \begin{pmatrix} b - c(2S^* + R^* + I^*) - d - \beta V^* & \delta' - cS^* & -cS^* & -\beta S^* \\ -cR^* & b' - c(S^* + 2R^* + I^*) - d - \delta' & \delta_i - cR^* & 0 \\ \beta V^* & 0 & -(\alpha + \alpha' + d + \delta_i) & \beta S^* \\ -\beta V^* & -\beta' V^* & b_v \alpha & -d_v - \beta S^* - \beta' R^* \end{pmatrix}$$

where $b' = b(1-\zeta)$, and $S^*$, $R^*$, $I^*$, $V^*$ stand for the number of (susceptible, resistant and infected) phytoplanktonic cells and virions (at any of the three fixed points evocated above).

*2.1. Stability analysis of the trivial equilibrium.* At the TE(0, 0, 0, 0), the Jacobian matrix associated with the SRIV-IPP model considering an extracellular resistance becomes:

$$J_1(0,0,0,0) = \begin{pmatrix} b-d & \delta' & 0 & 0 \\ 0 & b'-d-\delta' & 0 & 0 \\ 0 & 0 & -(\alpha+\alpha'+d) & 0 \\ 0 & 0 & b_v\alpha & -d_v \end{pmatrix}$$

and the Jacobian matrix associated with the SRIV-IPP model with an intracellular resistance is given by:

$$J_2(0,0,0,0) = \begin{pmatrix} b-d & \delta' & 0 & 0 \\ 0 & b'-d-\delta' & \delta_{\mathrm{i}} & 0 \\ 0 & 0 & -(\alpha+\alpha'+d+\delta_{\mathrm{i}}) & 0 \\ 0 & 0 & b_v\alpha & -d_v \end{pmatrix}$$

Since $J_1(0,0,0,0)$ and $J_2(0,0,0,0)$ are block diagonal and block triangular matrices, their eigenvalues correspond to those of their two $2 \times 2$ diagonal submatrices. According to the Routh-Hurwitz criteria, the eigenvalues of those submatrices have negative real parts whenever their trace is negative and their determinant is positive (Otto and Day 2011).

It is straightforward to show that the lower diagonal submatrices of $J_1(0,0,0,0)$ and $J_2(0,0,0,0)$ satisfy those criteria, so that the stability of the trivial equilibrium only depends on the eigenvalues of their upper diagonal submatrices. Interestingly, the upper diagonal submatrices of $J_1(0,0,0,0)$ and $J_2(0,0,0,0)$ are identical and since they are upper triangular, their eigenvalues are the diagonal entries, which must be negative for the TE to be stable. It is straightforward to show that the condition for the dominant eigenvalue to be $b-d$ is $b > -\frac{\delta'}{\zeta}$, which is obviously always hold. The condition for the TE to be stable therefore is;

$$\frac{b}{d} < 1 \tag{S28}$$

Accordingly, the intrinsic growth rate of a phytoplankton population ($\mathrm{R_0^H}$) appearing in table 2 corresponds to the left-hand side of S28.

*2.2. Stability analysis of the disease-free equilibrium.* At the DFE $(\mathrm{S}^*, 0,0,0)$, the expression of $\mathrm{S}^*$ simply reads

$$\mathrm{S}^* = \frac{b-d}{c} = \mathrm{K_N} \tag{S29}$$

and the general expressions of the Jacobian matrix associated with the SRIV-IPP model with an extracellular resistance simplify to:

$$J_1(\mathrm{S}^*,0,0,0)=\begin{pmatrix}-(b-d) & \delta'-(b-d) & -(b-d) & -\beta\mathrm{K_N}\\ 0 & -b\zeta-\delta' & 0 & \delta_{\mathrm{e}}\beta\mathrm{K_N}\\ 0 & 0 & -(\alpha+\alpha'+d) & (1-\delta_{\mathrm{e}})\beta\mathrm{K_N}\\ 0 & 0 & b_v\alpha & -d_v-(1-\delta_{\mathrm{e}})\beta\mathrm{K_N}\end{pmatrix}$$

whereas the one associated with the SRIV-IPP model with an intracellular resistance simplify to:

$$J_2(\mathrm{S}^*,0,0,0)=\begin{pmatrix}-(b-d) & \delta'-(b-d) & -(b-d) & -\beta\mathrm{K_N}\\ 0 & -b\zeta-\delta' & \delta_{\mathrm{i}} & 0\\ 0 & 0 & -(\alpha+\alpha'+d+\delta_{\mathrm{i}}) & \beta\mathrm{K_N}\\ 0 & 0 & b_v\alpha & -d_v-\beta\mathrm{K_N}\end{pmatrix}$$

Since $J_1(\mathrm{S}^*,0,0,0)$ and $J_2(\mathrm{S}^*,0,0,0)$ are block triangular matrices, their eigenvalues correspond to those of their two $2\times 2$ diagonal submatrices.

The upper diagonal submatrices of $J_1(\mathrm{S}^*,0,0,0)$ and $J_2(\mathrm{S}^*,0,0,0)$ are identical and their eigenvalues are the diagonal entries, which must be negative for the DFE to be stable. Given than $b > d$ for the phytoplankton population to grow, it is straightforward to show that the Routh-Hurwitz criteria are satisfied for those submatrices. Accordingly, the stability of the DFE ($\mathrm{S}^*$,0,0,0) only depends on the eigenvalues of the lower diagonal submatrices of the above Jacobians. Although the traces of those submatrices are obviously negative, their determinants must be positive for the DFE to be stable, which led to the following condition when considering an extracellular resistance;

$$(1-\delta_{\mathrm{e}})\beta\mathrm{K_N}\left(\frac{\alpha}{(\alpha+\alpha'+d)}b_v-1\right)\frac{1}{d_v}<1 \qquad \text{(S30)}$$

and to a similar condition when resistance is assumed to be intracellular;

$$\beta K_N \left(\frac{\alpha}{(\alpha + \alpha' + d + \delta_i)} b_v - 1\right) \frac{1}{d_v} < 1 \tag{S31}$$

Albeit slightly different, those conditions carry the same rational; whenever they are lower than 1, the DFE is stable since the virus is unable to spread into a population made of susceptible phytoplankton at their demographic equilibrium. Conversely, when such a condition is not verified, one expects the virus to be able to spread by infecting the susceptible individuals. Accordingly, the left-hand side of S30 and S31 represent the initial growth rate of the viral population, i.e. the virus basic reproduction number $\left(R_0^V\right)$ appearing in table 3.

*2.3. Stability analysis of the disease equilibrium*. At the DE ($S^*$, $R^*$, $I^*$, $V^*$), the formal expression of the densities of individuals can still be identified for each of the two SRIV-IPP models, although this requires the convoluted calculations described in Supporting Information, Heading 2. By applying this analytical procedure to equations S26.*a*, S26.*c*, S26.*d*, we first expressed $I^*$ and $V^*$ as functions of $R^*$ and then found the expression of $S^*$ for the SRIV-IPP model considering an extracellular resistance:

$$I^*(R^*) = A_1 S^* V^*(R^*) \tag{S32.$a$}$$

$$S^* = A_2 \tag{S32.$b$}$$

$$V^*(R^*) = \frac{A_3 + B_3 R^*}{(\beta + c A_1 A_2) A_2} \tag{S32.$c$}$$

where $A_1 = \frac{(1-\delta_e)\beta}{(\alpha+\alpha'+d)}$, $A_2 = \frac{d_v}{(b_v \alpha A_1 - (1-\delta_e)\beta)}$, $A_3 = (b - d - cA_2)A_2$ and $B_3 = \delta' - cA_2$.

The three functions above were then substituted for $S^*$, $I^*$ and $V^*$ in equation S26.*b*, to derive a quadratic equation allowing for the identification of R* for the first IPP model:

$$f_1(R^*) = A_4 + B_4 R^* + C_4 R^{*2} = 0 \tag{S33}$$

Where $A_4 = \beta\delta_e(b - d - cA_2)A_2$, $B_4 = \beta\delta_e(p - cA_2) + \beta(b' - d - cA_2 - \delta') - c\delta' A_1 A_2 - cb\zeta A_1 A_2$ and $C_4 = -c(\beta + \delta' A_1)$

As $C_4$ is negative, this function corresponds to a concave parabola. Given that the condition for $A_2$ to be positive is that the $R_0^V > 1$ (S30), which is the case to reach the DE, the equation S33 has only one positive solution, so that the expression of $R^*$ is given by:

$$R^* = \frac{-B_4 - \sqrt{B_4^2 - 4A_4C_4}}{2C_4} \tag{S34}$$

Equation S34 was subsequently used to identify $I^*$ and $V^*$ according to the relationships S32.*a* and S32.*c*.

The same method was applied to equations S27.*a*, S27.*c*, S27.*d* to express $S^*$, $I^*$ and $V^*$ as functions of $R^*$ for the SRIV-IPP model considering an intracellular resistance, which led to:

$$I^*(R^*) = A_1 S^*(R^*) V^*(R^*) \tag{S35.a}$$

$$S^*(R^*) = A_2 + B_2 R^* \tag{S35.b}$$

$$V^*(R^*) = \frac{A_3 + B_3 R^*}{\beta + cA_1(A_2 + B_2 R^*)} \tag{S35.c}$$

where $A_1 = \frac{\beta}{(\alpha + \alpha' + d + \delta_i)}$, $A_2 = \frac{d_v}{(b_v \alpha A_1 - \beta)}$, $B_2 = \frac{\beta'}{(b_v \alpha A_1 - \beta)}$, $A_3 = b - d - cA_2$ and $B_3 = \frac{\delta'}{(A_2 + B_2 R^*)} - c(B_2 + 1)$.

Combining those three functions and equation S27.*b* provided a quadratic equation allowing for the identification of $R^*$ for the second SRIV-IPP model:

$$f_2(R^*) = A_4 + B_4 R^* + C_4 R^{*2} = 0 \tag{S36}$$

Where $A_4 = \delta_i(b - d - cA_2)A_1A_2$, $B_4 = \beta(b' - d - cA_2 - \delta') + (b - d - 2cA_2)\delta_iA_1B_2 + \delta_i(\delta' - cA_2)A_1 - c(b\zeta - \delta')A_1A_2$ and $C_4 = -c(b\zeta - \delta')A_1B_2 - c\beta(B_2 + 1) - c\delta'A_1 - c\delta_iA_1B_2(B_2 + 1)$.

It is straightforward to show that $C_4$ is negative and, following the same logic as above, that $A_4 > 0$ if $R_0^V > 1$ (S31). The equilibrium level of $R^*$ for the second IPP model was then set to correspond to the only positive solution of the above quadratic equation (that has the same expression as S34), and subsequently used to identify $S^*$, $I^*$ and $V^*$ according to the relationships S35.*a* to S35.*c*.

The conditions for the DE to be stable can be found using the following Routh–Hurwitz criteria (Otto and Day 2011):

$$a_1 > 0 \qquad (S37.a)$$

$$a_3 > 0 \qquad (S37.b)$$

$$a_4 > 0 \qquad (S37.c)$$

$$a_1a_2a_3 > a_3^2 + a_1^2a_4 \qquad (S37.d)$$

where $a_1$ to $a_4$ denote the coefficients of the characteristic polynomial of the Jacobian:

$$\lambda^4 + a_1\lambda^3 + a_2\lambda^2 + a_3\lambda + a_4 \qquad (S38)$$

For the Jacobian matrix associated with the SRIV-IPP model considering an extracellular resistance, these coefficients are given by:

$$a_1 = (\alpha + \alpha' + d) + \beta(1 - \delta_e)b_v\frac{\alpha}{(\alpha + \alpha' + d)}S^* + cS^* + cR^* + \beta\delta_e\frac{S^*V^*}{R^*} + \delta'\frac{R^*}{S^*}$$

$$a_2 = (\alpha + \alpha' + d)\left(cS^* + cR^* + \beta\delta_e\frac{S^*V^*}{R^*} + \delta'\frac{R^*}{S^*}\right) + c\beta S^*V^* + c\delta'R^* + c\beta\delta_e\frac{S^{*2}V^*}{R^*}$$

$$+\beta b_v \frac{\alpha}{(\alpha+\alpha'+d)}(1-\delta_{\mathrm{e}})\left(c\mathrm{S}^{*2}+c\mathrm{S}^*\mathrm{R}^*+\beta\delta_{\mathrm{e}}\frac{\mathrm{S}^{*2}\mathrm{V}^*}{\mathrm{R}^*}+\delta'\mathrm{R}^*\right)+c\delta'\frac{\mathrm{R}^{*2}}{\mathrm{S}^*}-\beta^2(1-\delta_{\mathrm{e}})\mathrm{S}^*\mathrm{V}^*$$

$$a_3 = c(\alpha+\alpha'+d)\left(\beta\delta_{\mathrm{e}}\mathrm{S}^*\mathrm{V}^*+\delta'\mathrm{R}^*+\beta\delta_{\mathrm{e}}\frac{\mathrm{S}^{*2}\mathrm{V}^*}{\mathrm{R}^*}+\delta'\frac{\mathrm{R}^{*2}}{\mathrm{S}^*}\right)+ c\beta\delta'(1-\delta)\mathrm{R}^*\mathrm{V}^*$$

$$+\beta^2\delta_{\mathrm{e}}\delta'(1-\delta)\mathrm{S}^*\mathrm{V}^*+c\beta^2\delta_{\mathrm{e}}(1-\delta_{\mathrm{e}})\frac{\mathrm{S}^{*2}\mathrm{V}^{*2}}{\mathrm{R}^*}+\beta^2(1-\delta_{\mathrm{e}})b_v\alpha\mathrm{S}^*\mathrm{V}^*$$

$$+c\beta b_v\frac{\alpha}{(\alpha+\alpha'+d)}(1-\delta_{\mathrm{e}})\left(\beta\mathrm{S}^{*2}\mathrm{V}^*+\beta\delta_{\mathrm{e}}\frac{\mathrm{S}^{*3}\mathrm{V}^*}{\mathrm{R}^*}+\delta'\mathrm{S}^*\mathrm{R}^*+\delta'\mathrm{R}^{*2}\right)$$

$$-\beta^3\delta_{\mathrm{e}}(1-\delta_{\mathrm{e}})\frac{\mathrm{S}^{*2}\mathrm{V}^{*2}}{\mathrm{R}^*}-c\beta^2(1-\delta_{\mathrm{e}})\mathrm{S}^*\mathrm{R}^*\mathrm{V}^*-c\beta^2(1-\delta_{\mathrm{e}})\mathrm{S}^{*2}\mathrm{V}^*$$

$$-\beta^2(\alpha+\alpha'+d)(1-\delta_{\mathrm{e}})\mathrm{S}^*\mathrm{V}^*$$

$$a_4 = \beta^3\delta_{\mathrm{e}}(1-\delta_{\mathrm{e}})b_v\alpha\frac{\mathrm{S}^{*2}\mathrm{V}^{*2}}{\mathrm{R}^*}+c\beta^3 b_v\frac{\alpha}{(\alpha+\alpha'+d)}\delta_{\mathrm{e}}(1-\delta_{\mathrm{e}})^2\frac{\mathrm{S}^{*3}\mathrm{V}^{*2}}{\mathrm{R}^*}+c\beta^2\delta_{\mathrm{e}}(1-\delta_{\mathrm{e}})b_v\alpha\mathrm{S}^{*2}\mathrm{V}^*$$

$$+c\beta^2(1-\delta_{\mathrm{e}})b_v\alpha\mathrm{S}^*\mathrm{R}^*\mathrm{V}^*+c\beta^2 b_v\frac{\alpha}{(\alpha+\alpha'+d)}\delta'(1-\delta_{\mathrm{e}})^2\mathrm{S}^*\mathrm{R}^*\mathrm{V}^*$$

$$+\beta^2(\alpha+\alpha'+d)\delta_{\mathrm{e}}\delta'(1-\delta_{\mathrm{e}})\mathrm{S}^*\mathrm{V}^*-\beta^3(\alpha+\alpha'+d)\delta_{\mathrm{e}}(1-\delta_{\mathrm{e}})\frac{\mathrm{S}^{*2}\mathrm{V}^{*2}}{\mathrm{R}^*}$$

$$-c\beta^3\delta_{\mathrm{e}}(1-\delta_{\mathrm{e}})^2\frac{\mathrm{S}^{*3}\mathrm{V}^{*2}}{\mathrm{R}^*}-\beta^2\delta_{\mathrm{e}}\delta'(1-\delta_{\mathrm{e}})b_v\alpha\mathrm{S}^*\mathrm{V}^*-c\beta^2(\alpha+\alpha'+d)(1-\delta_{\mathrm{e}})\mathrm{S}^*\mathrm{R}^*\mathrm{V}^*$$

$$-c\beta^2\delta'(1-\delta_{\mathrm{e}})^2\mathrm{S}^*\mathrm{R}^*\mathrm{V}^*-c\beta^2(\alpha+\alpha'+d)\delta_{\mathrm{e}}(1-\delta_{\mathrm{e}})\mathrm{S}^{*2}\mathrm{V}^*$$

while for the Jacobian matrix associated with the SRIV-IPP model with intracellular resistance, they can be shown to be;

$$a_1 = (\alpha+\alpha'+d+\delta_{\mathrm{i}})+\beta b_v\frac{\alpha}{(\alpha+\alpha'+d+\delta)}\mathrm{S}^*+c\mathrm{S}^*+c\mathrm{R}^*+\delta'\frac{\mathrm{R}^*}{\mathrm{S}^*}+\delta_{\mathrm{i}}\frac{\mathrm{I}^*}{\mathrm{R}^*}$$

$$a_2 = (\alpha + \alpha' + d + \delta_{\mathrm{i}})\left(c\mathrm{S}^* + c\mathrm{R}^* + \delta_{\mathrm{i}}\frac{\mathrm{I}^*}{\mathrm{R}^*} + \delta'\frac{\mathrm{R}^*}{\mathrm{S}^*}\right) + c\beta\mathrm{S}^*\mathrm{V}^* + c\delta'\mathrm{R}^*$$

$$+\beta b_v\frac{\alpha}{(\alpha + \alpha' + d + \delta_{\mathrm{i}})}\left(c\mathrm{S}^{*2} + c\mathrm{S}^*\mathrm{R}^* + \delta'\mathrm{R}^* + \delta_{\mathrm{i}}\frac{\mathrm{S}^*\mathrm{I}^*}{\mathrm{R}^*}\right) + \delta_{\mathrm{i}}\delta'\frac{\mathrm{I}^*}{\mathrm{S}^*} + c\delta_{\mathrm{i}}\frac{\mathrm{S}^*\mathrm{I}^*}{\mathrm{R}^*}$$

$$+c\delta'\frac{\mathrm{R}^{*2}}{\mathrm{S}^*} - \beta^2\mathrm{S}^*\mathrm{V}^*$$

$$a_3 = (\alpha + \alpha' + d + \delta_{\mathrm{i}})\left(\delta_{\mathrm{i}}\delta'\frac{\mathrm{I}^*}{\mathrm{S}^*} + c\delta'\mathrm{R}^* + c\delta'\frac{\mathrm{R}^{*2}}{\mathrm{S}^*} + c\delta_{\mathrm{i}}\frac{\mathrm{S}^*\mathrm{I}^*}{\mathrm{R}^*}\right) + c\beta\delta'\mathrm{R}^*\mathrm{V}^* + c\beta\delta_{\mathrm{i}}\mathrm{S}^*\mathrm{V}^*$$

$$+c\beta\delta_{\mathrm{i}}\frac{\mathrm{S}^*\mathrm{I}^*\mathrm{V}^*}{\mathrm{R}^*} + \beta b_v\frac{\alpha}{(\alpha + \alpha' + d + \delta_{\mathrm{i}})}\left(\delta_{\mathrm{i}}\delta'\mathrm{I}^* + c\beta\mathrm{S}^{*2}\mathrm{V}^* + c\delta'\mathrm{S}^*\mathrm{R}^* + c\delta'\mathrm{R}^{*2} + c\delta_{\mathrm{i}}\frac{\mathrm{S}^{*2}\mathrm{I}^*}{\mathrm{R}^*}\right)$$

$$+\beta^2 b_v\alpha\mathrm{S}^*\mathrm{V}^* + \beta\beta'\delta_{\mathrm{i}}\mathrm{S}^*\mathrm{V}^* - \beta^2(\alpha + \alpha' + d + \delta_{\mathrm{i}})\mathrm{S}^*\mathrm{V}^* - c\beta^2\mathrm{S}^{*2}\mathrm{V}^* - c\beta^2\mathrm{S}^*\mathrm{R}^*\mathrm{V}^* - \beta\delta_{\mathrm{i}}\delta'\mathrm{V}^*$$

$$-\beta^2\delta_{\mathrm{i}}\frac{\mathrm{S}^*\mathrm{I}^*\mathrm{V}^*}{\mathrm{R}^*}$$

$$a_4 = c\beta\beta'(\alpha + \alpha' + d + \delta_{\mathrm{i}})\mathrm{S}^*\mathrm{R}^*\mathrm{V}^* + c\beta^2 b_v\frac{\alpha}{(\alpha + \alpha' + d + \delta_{\mathrm{i}})}\left(\delta_{\mathrm{i}}\frac{\mathrm{S}^{*2}\mathrm{I}^*\mathrm{V}^*}{\mathrm{R}^*} + \delta_{\mathrm{i}}\mathrm{S}^{*2}\mathrm{V}^* + \delta'\mathrm{S}^*\mathrm{R}^*\mathrm{V}^*\right)$$

$$+c\beta^2\beta'\mathrm{S}^*\mathrm{R}^*\mathrm{V}^{*2} + c\beta^2 b_v\alpha\mathrm{S}^*\mathrm{R}^*\mathrm{V}^* + \beta^2\delta_{\mathrm{i}}\delta'\mathrm{S}^*\mathrm{V}^* + \beta\beta'\delta_{\mathrm{i}}\delta'\mathrm{R}^*\mathrm{V}^* + c\beta\beta'\delta_{\mathrm{i}}\mathrm{S}^{*2}\mathrm{V}^*$$

$$+\beta^2 b_v\alpha\delta_{\mathrm{i}}\frac{\mathrm{S}^*\mathrm{I}^*\mathrm{V}^*}{\mathrm{R}^*} - c\beta^2(\alpha + \alpha' + d + \delta_{\mathrm{i}})\mathrm{S}^*\mathrm{R}^*\mathrm{V}^* - \beta^2(\alpha + \alpha' + d + \delta_{\mathrm{i}})\delta_{\mathrm{i}}\frac{\mathrm{S}^*\mathrm{I}^*\mathrm{V}^*}{\mathrm{R}^*}$$

$$-\beta^2 b_v\frac{\alpha}{(\alpha + \alpha' + d + \delta_{\mathrm{i}})}\delta_{\mathrm{i}}\delta'\mathrm{S}^*\mathrm{V}^* - c\beta^2\delta'\mathrm{S}^*\mathrm{R}^*\mathrm{V}^* - \beta^2\beta'\delta_{\mathrm{i}}\mathrm{S}^*\mathrm{V}^{*2} - c\beta^2\delta_{\mathrm{i}}\mathrm{S}^{*2}\mathrm{V}^*$$

$$-c\beta\beta'\delta'\mathrm{R}^{*2}\mathrm{V}^* - c\beta^2\delta_{\mathrm{i}}\frac{\mathrm{S}^{*2}\mathrm{I}^*\mathrm{V}^*}{\mathrm{R}^*}$$

The above expressions for both SRIV-IPP models were used, together with the expressions of $S^*$, $R^*$, $I^*$, and $V^*$, derived when resistance is extracellular (S32*a-c* and S34 with $A_4$,$B_4$ and $C_4$ defined from S33) and intracellular (S35*a-c* and S34 with $A_4$,$B_4$ and $C_4$ defined from S36), were then used to numerically assess the local stability of the DE in both of these cases.

**Heading 5. Estimation of the encounter rate between phytoplankton and virus $\beta$.**

A widely used expression to estimate the encounter rate $(\beta)$ between a unicellular host and its virus in a marine environment was first proposed and used by Murray and Jackson (1992, (Murray and Jackson 1992));

$$\beta = Sh.2.\pi.D_h.\frac{k.\mathrm{T}}{3.\pi.\eta.D_v} \tag{S1}$$

This relationships links the desired encounter rate with the host ($D_h$) and virus ($D_v$) diameters, and standard parameters describing the abiotic environment; the Boltzmann constant ($k$), the water viscosity ($\eta$) - depending on both temperature (T) and salinity - and the Sherwood number ($Sh$) that accounts for the motility of the particles (increasing with motility from 1 for non-motile species). To be consistent with the (experimental) conditions that allowed other model parameters (See table 1 in the main text) to be estimated, the temperature and salinity were set to 293.15 K (20°C) and 38.57 psu (Aydogdu et al. 2023), which provided the value of T and $\eta$. Meanwhile, we set $D_v$ to the mean virus capsid diameter of OtV estimated to be equal to 122 [113 ; 131] nm (Derelle et al. 2008).

To derive an estimate of $\beta$ from S1, the only quantity left to quantify was therefore the diameter of *O. tauri* ($D_h$) and we aimed at using the Equivalent Spherical Diameter (ESD) as a typical estimate of $D_h$ (Murray and Jackson 1992).

To do so, we estimated the mean and variance of the length ($l$) and width ($w$) of *O. tauri* cells from measures made on 116 individuals (Chrétiennot-Dinet et al. 1995). Assuming, as described for unicellular eukaryotes (Giometto et al. 2013), that those cell's dimension are log-normally distributed, we calculated confidence interval for the average $l$ (0.97 µm [0.69 ; 1.25]) and $w$ (0.70 µm [0.53 ; 0.87]). We then used the two log-normal distributions to sample a total of $10^7$ length and width of *O. tauri* and used them to calculate $10^7$ cell volumes considering the cell shape as a rotational ellipsoid (i.e. according to $V_{re} = \frac{\pi}{6} l w^2$, (Olenina et al. 2006)). These 'initial' volumes were subsequently assimilated to spherical volumes in order to derive the $10^7$ corresponding ESD, from which we calculated the mean and its 95% confidence interval (0.7791 µm [0.7786 ; 0.7799]). Ultimately, we applied S1 to this distribution of ESD in order to obtain a distribution of encounter rates, from which we obtain the mean estimate of $\beta$ equals to $1.37\times10^{-6}$ $ml.d^{-1}$ that appear in table 1 in the main text.

**Heading 6. Test of the accuracy of the approximate expression of $R^*$ at the Disease Equilibrium for the SRIV-GM model of phytoplankton-virus interaction.**

To check the accuracy of the approximate expression of $R^*$ obtained by Taylor expansion at the Disease Equilibrium (equations S10.*a*-*c*), we compared it with the non-approximated value of $R^*$ that we evaluated by numerically solving the cubic equation (S9) derived in supporting text Heading 2 (by using the function "polyroot" of the R Base package, R version 4.2.0). The tables below summarise the outcomes of those comparisons achieved for the entire range of probability of mutation ($\mu$) considered in this contribution and under the hypotheses that resistance is extracellular (table S2) or intracellular (table S3).

**Table S2. Summary of the non-approximate (numerical) and approximate values of the abundance of extracellular resistant hosts $R^*$.** a. In the absence of PCD. b. In the presence of PCD (with $\alpha' = 2\alpha$).

(a)

| Mutation event ($\mu$) | Numeric | Approximate | Δ% |
|---|---|---|---|
| $10^{-9}$ | 1412409 | 1412409 | 0 |
| $10^{-8}$ | 1412409 | 1412409 | 0 |
| $10^{-7}$ | 1412409 | 1412409 | 0 |
| $10^{-6}$ | 1412409 | 1412409 | 0 |
| $10^{-5}$ | 1412409 | 1412409 | 0 |
| $10^{-4}$ | 1412409 | 1412409 | 0 |
| $10^{-3}$ | 1412409 | 1412409 | 0 |
| $10^{-2}$ | 1412409 | 1412409 | 0 |

(b)

| Mutation event ($\mu$) | Numeric | Approximate | Δ% |
|---|---|---|---|
| $10^{-9}$ | 1411718 | 1411718 | 0 |
| $10^{-8}$ | 1411718 | 1411718 | 0 |
| $10^{-7}$ | 1411718 | 1411718 | 0 |
| $10^{-6}$ | 1411718 | 1411718 | 0 |
| $10^{-5}$ | 1411718 | 1411718 | 0 |
| $10^{-4}$ | 1411718 | 1411718 | 0 |
| $10^{-3}$ | 1411718 | 1411718 | 0 |
| $10^{-2}$ | 1411718 | 1411718 | 0 |

**Table S3. Summary of the non-approximate (numerical) and approximate values of the abundance of intracellular resistant hosts R*.** a. In the absence of PCD. b. In the presence of PCD (with $\alpha' = 2\alpha$).

(a)

| **Mutation event ($\mu$)** | **Numeric** | **Approximate** | **Δ%** |
|---|---|---|---|
| $10^{-9}$ | 1354617 | 1354617 | 0 |
| $10^{-8}$ | 1354617 | 1354617 | 0 |
| $10^{-7}$ | 1354617 | 1354617 | 0 |
| $10^{-6}$ | 1354617 | 1354617 | 0 |
| $10^{-5}$ | 1354617 | 1354617 | 0 |
| $10^{-4}$ | 1354618 | 1354618 | 0 |
| $10^{-3}$ | 1354623 | 1354623 | 0 |
| $10^{-2}$ | 1354678 | 1354678 | 0 |

(b)

| **Mutation event ($\mu$)** | **Numeric** | **Approximate** | **Δ%** |
|---|---|---|---|
| $10^{-9}$ | 1241030 | 1241030 | 0 |
| $10^{-8}$ | 1241030 | 1241030 | 0 |
| $10^{-7}$ | 1241030 | 1241030 | 0 |
| $10^{-6}$ | 1241030 | 1241030 | 0 |
| $10^{-5}$ | 1241031 | 1241031 | 0 |
| $10^{-4}$ | 1241032 | 1241032 | 0 |
| $10^{-3}$ | 1241046 | 1241046 | 0 |
| $10^{-2}$ | 1241183 | 1241184 | $1.69\times10^{-5}$ |

#### Supporting Figures

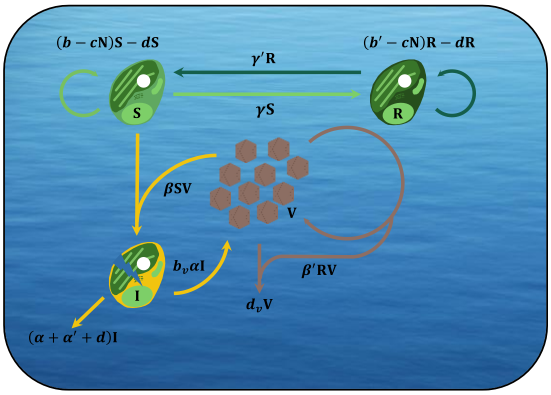


**(a) Random Phenotypic Plasticity (RPP)**

$$\frac{dS}{dt} = (b - cN)S - dS + \gamma' R - \gamma S - \beta SV$$

$$\frac{dR}{dt} = (b' - cN)R - dR + \gamma S - \gamma' R$$

$$\frac{dI}{dt} = \beta SV - (\alpha + \alpha' + d)I$$

$$\frac{dV}{dt} = b_v \alpha I - d_v V - \beta SV - \beta' RV$$

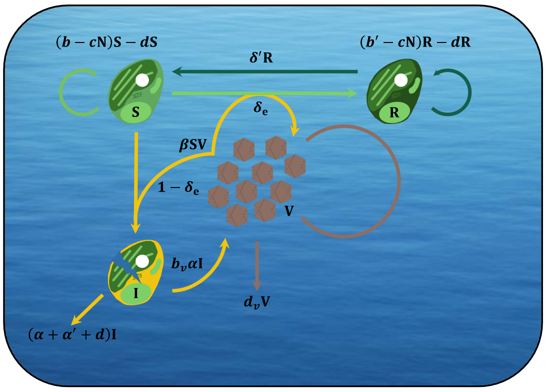


**(b) Induced Phenotypic Plasticity (IPP) – Ext.**

$$\frac{dS}{dt} = (b - cN)S - dS + \delta' R - \beta SV$$

$$\frac{dR}{dt} = (b' - cN)R - dR + \delta_e \beta SV - \delta' R$$

$$\frac{dI}{dt} = (1 - \delta_e)\beta SV - (\alpha + \alpha' + d)I$$

$$\frac{dV}{dt} = b_v \alpha I - d_v V - (1 - \delta_e)\beta SV$$

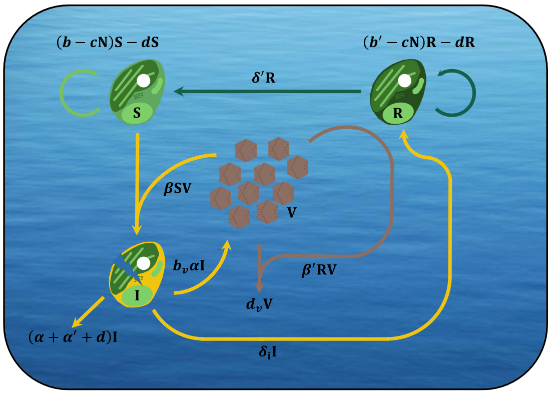


**(c) Induced Phenotypic Plasticity (IPP) – Int.**

$$\frac{dS}{dt} = (b - cN)S - dS + \delta' R - \beta SV$$

$$\frac{dR}{dt} = (b' - cN)R - dR + \delta_i I - \delta' R$$

$$\frac{dI}{dt} = \beta SV - (\alpha + \alpha' + d + \delta_i)I$$

$$\frac{dV}{dt} = b_v \alpha I - d_v V - \beta SV - \beta' RV$$

**Fig S1. Summary of SRIV models of phytoplankton-virus interaction when antiviral resistance is associated to phenotypic plasticity.** (a) Random Phenotypic Plasticity (SRIV-RPP). Extracellular and intracellular mechanisms of resistance can be represented using the set of equations 2.8 to 2.11 in the main text (and reproduced below) by setting $\beta' = 0$ and $\beta' = \beta$, respectively. (b) Virus-Induced Phenotypic Plasticity (SRIV-IPP) when resistance is extracellular. A contact between a susceptible cell and a virion allows for the cell to become resistant ($\delta_e$) or infected ($1 - \delta_e$). This can be described by using equations 2.12 to 2.15 in the main text while setting $\delta_i$=0, which lead to the system of equations below. (c) Virus-Induced Phenotypic Plasticity (SRIV-IPP) when resistance is intracellular. An infected cell can become resistant by clearing and switching ($\delta_i$). The same set of equations, i.e. 2.12 to 2.15 in the main text, can be used while considering $\delta_e$=0, which lead to the model appearing below.

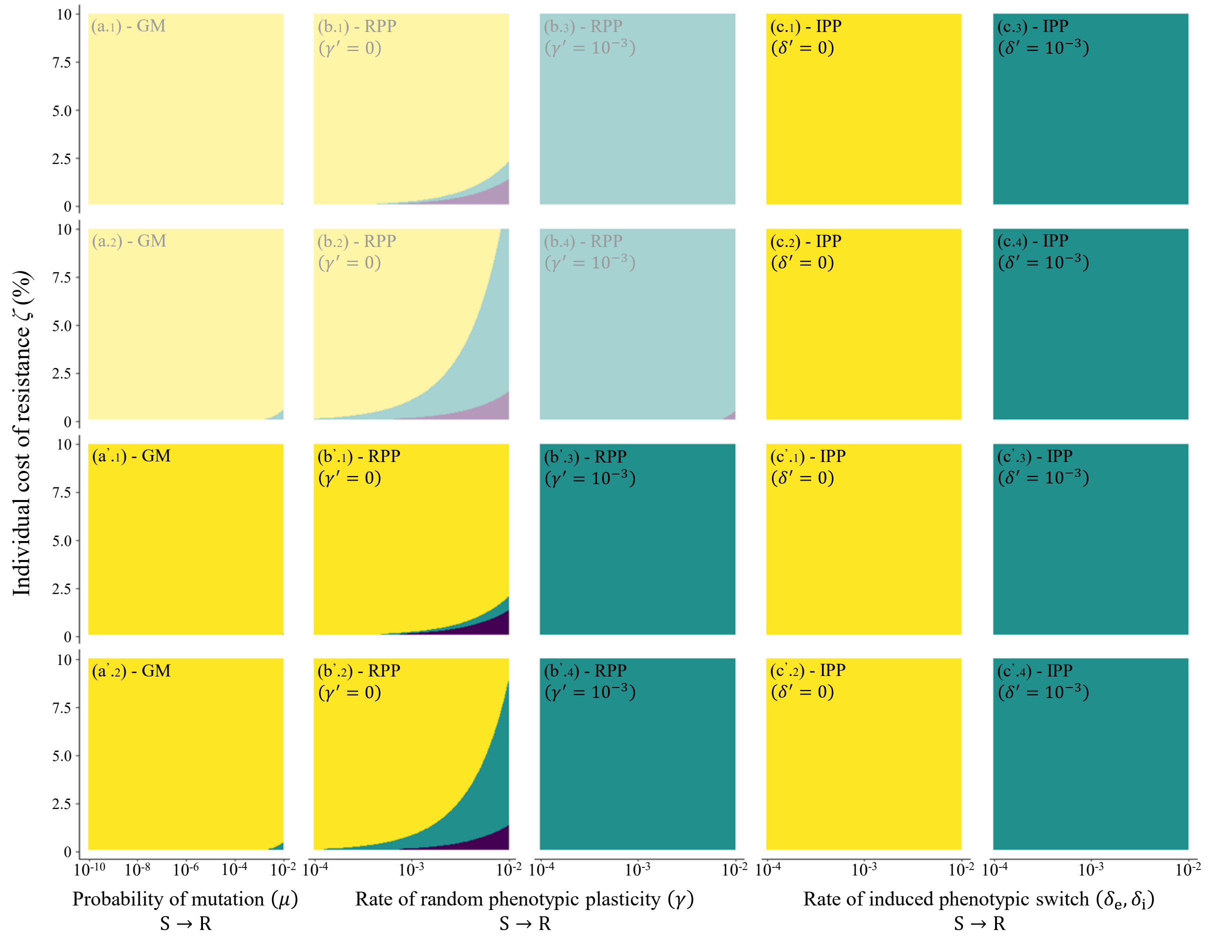


**Figure S2. Effects of antiviral resistance on the long-term dynamics of *O. tauri*–OtV interaction**. (a-a') Antiviral resistance associated to Genomic Mutations (SRIV-GM model). (b-b') Antiviral resistance acquired through Random Phenotypic Plasticity (SRIV-RPP model) with a rate of phenotypic switch from R to S ($\gamma'$) equals to zero (b-b'.1-2) and $10^{-3}$ (b-b'.3-4). (c-c') Antiviral resistance acquired through virus Induced Phenotypic Plasticity (SRIV-IPP model) with a rate of phenotypic switch from R to S ($\delta'$) equals to zero (c-c'.1-2) and $10^{-3}$ (c-c' .3-4). a, b and c show the effect of antiviral resistance when it corresponds to intracellular mechanism (as described in the main text), while a', b' and c' illustrate that the predicted outcomes of the *O. tauri*-OtV interaction are highly similar when resistance is extracellular. For a-a', b-b' and c-c', those outcomes are shown in the absence (panels 1 and 3) or in the presence (panels 2 and 4) of PCD (with $\alpha' = 2\alpha$). In each panel, yellow and green areas correspond to *O. tauri*–OtV coexistence through oscillatory and stable dynamics, respectively, while purple areas show the conditions where the OtV is unable to spread. Figures S1a.1-2 and S1b.1-4 appearing in pastel colours are those described in the main text.
